# Preservation of dopamine neurotransmission during nigrostriatal neuron loss in rat Parkinson’s model: evidence for increased dopamine signaling in substantia nigra

**DOI:** 10.1101/2025.09.12.675909

**Authors:** Ashley Galfano, Robert McManus, Walter Navarrete, Sampada Chaudhari, Christopher Bishop, Michael F. Salvatore

## Abstract

During progressive nigrostriatal neuron loss in Parkinson’s disease (PD), compensatory mechanisms are thought to maintain dopamine (DA) signaling at levels sufficient to mitigate locomotor impairment prior to substantial neuron loss. Whereas increased DA turnover in striatum has been considered a keystone compensatory mechanism indicative of augmented DA signaling, recent work indicates that increased DA biosynthesis in substantia nigra (SN), not striatum, compensates for tyrosine hydroxylase (TH) protein and neuronal loss. To extend interrogation of compensatory mechanisms that augment DA signaling during nigrostriatal neuron loss, we contemporaneously evaluated extracellular DA against tissue DA levels and quantified TH protein in the striatum and SN. Our unilateral 6-hydroxydopamine (6-OHDA) lesion approach produces progressive neuronal loss between 7 and 28 days, enabling evaluation of multiple DA signaling components in striatum vs SN. Loss of TH was ∼90% in striatum and ∼70% in the SN by 7 days after lesion induction. However, whereas loss of tissue DA matched TH loss in striatum (>90%) on both days after lesion, tissue DA loss in SN occurred only at day 28 (36%), despite major TH loss by day 7. This preservation of nigral DA tissue levels during lesion progression was associated with increased extracellular DA in SN after K+-dependent depolarization in striatum, which was not evident in a sham-operation group at the same time, early after lesion induction, signifying a DA lesion-specific adaptation in SN. In contrast in the striatum, lesion abolished the robust increase in extracellular DA, as seen in the sham-operation group. Together, these results indicate compensatory mechanisms that augment nigrostriatal DA signaling are engaged in the SN, not striatum, during progressive nigrostriatal neuron loss.

## Introduction

Motor impairment in Parkinson’s disease (PD) first occurs when nigrostriatal dopamine (DA) neuron loss exceeds 50%, coincident with >70 to 80% loss of tyrosine hydroxylase (TH) in the striatum (Bernheimer et al., 1973; Bezard et al., 2001; Kordower et al., 2013). Well after PD diagnosis, 10-50% of neurons remain in the SN, whereas in the striatum, TH and DA transporter proteins are virtually undetectable (Kordower et al., 2013; Furukawa et al., 2022). Paradoxically, motor impairment still increases in severity even though striatal DA markers are undetectable (Furukawa et al., 2022; Kordower et al., 2013; Perlmutter et al., 2014; Tabbal et al., 2012). Therefore, in human PD, the metrics of striatal DA loss is not linear with motor impairment, making our understanding of how nigrostriatal DA signaling affects motor impairment far from complete.

Given that severe striatal TH loss is present at the onset of motor impairment, engagement of compensatory mechanisms in DA signaling have been proposed to maintain motor function despite nigrostriatal neuron loss (Blesa et al., 2012; 2017; Collier et al., 2007; Fearnley and Lees, 1991; Pifl and Hornykiewicz, 2006; Zigmond, 1997). However, current evidence does not point to a compensatory mechanism occurring in striatum to augment DA signaling. Increased striatal DA metabolism and/or turnover occurs only after onset of motor impairment, accompanied by severe TH protein loss in striatum (Bezard et al., 2001; Blesa et al, 2012; Pifl and Hornykiewicz, 2006; Kasanga et al., 2023; Salvatore, 2014; 2024). Increased DA biosynthesis, by increased TH phosphorylation as a compensatory mechanism, does not occur in striatum (Salvatore, 2014; Kasanga et al., 2023). While evidence supports that post-synaptic DA receptor expression can upregulate in response to DA depletion (Araki et al., 2000; Feyder et al., 2011), the increased expression of DA D1 receptors is not consistently substantiated in striatum (Kasanga et al., 2023). The lack of evidence for plasticity in striatum has led to the idea that non-dopaminergic mechanisms serve as possible compensatory processes, including increased basal ganglia circuit activity and engagement of circuits from the motor cortex or thalamic nuclei (Bezard et al., 1999; Blesa et al., 2017). Endogenous DA must still be available to drive basal ganglia circuitry. As such, others have reported that extracellular DA levels may increase in response to neuronal loss (Perez et al., 2008; Pérez-Taboada, et al., 2020; Cramb et al., 2023) to compensate against striatal TH protein and DA tissue loss (Kordower et al., 2013; Kasanga et al., 2023). However, it is currently unknown whether DA tissue loss at the magnitude seen at onset of motor impairment would directly limit DA release capacity in striatum.

Historically, the loss of striatal DA has been argued to be central to disruption of normal basal ganglia output that produces impoverished motor function. However, an oftenoverlooked component of DA signaling in the nigrostriatal pathway is also present in the SN, bearing the same regulation steps of DA signaling that are present in striatum (Cragg and Greenfield, 1997; Cragg et al., 1997; Geffen et al., 1976; Salvatore, 2024). As loss of DA markers within the SN occur at a slower rate compared to the striatum in human PD (Bezard et al., 2001; Kordower et al., 2013; Matuskey et al., 2020), it plausible that remaining cell bodies in SN could be a site of compensatory mechanisms to influence motor function. Indeed, it is plausible that somatodendritic release of DA in the SN (as shown in previous studies ((Geffen et al., 1976; Hoffman & Gerhardt, 1998; Gerhardt et al., 2002; Bergquist et al., 2003; Sarre et al., 1990, 1992, 2004) may drive basal ganglia output to influence motor function through local release in the SN (Robertson and Robertson, 1988, 1999; Robertson et al., 1991) and subsequent DA binding to post-synaptic DA D1 receptors (Kliem et al., 2007, 2010; rev. Salvatore, 2024).

Contralateral or inter-hemispheric compensation may also play a role in maintaining DA signaling and locomotion during striatal DA loss. In fact, the transition to Hoehn-Yahr stage 1 to 2 is characterized by progression from unilateral to bilateral motor impairment. Evidence in both preclinical and clinical literature suggests hemisphere asymmetry in neuronal processes which drive the motor symptom lateralization (Riederer et al., 2017; Li et al., 2020). For example, in unilaterally 6-hydroxydopamine (6-OHDA) lesioned rats, stimulating the contralateral SN increases ipsilateral striatal DA release, whereas stimulating the ipsilateral SN does not (Fox et al., 2016). Following unilateral 6-OHDA lesion, reorganization of inter-hemispheric pathways within higher cortical regions may also influence movement (Javor-Duray et al., 2017). Arguably, the unbalanced degeneration and dysfunction that characterizes PD, at least in early stages, may provide an avenue for therapeutic targeting (Slevin et al., 2005). However, it is not known exactly how, where, and extent to which tissue DA and TH protein loss during nigrostriatal neuron loss would affect extracellular DA availability. Understanding the nature of these inherent responses to maintain DA signaling in response to nigrostriatal neuron loss can reveal which intervention points would be most effective to improve motor function.

Using a unilateral 6-OHDA regimen that produces a progressive loss of nigrostriatal neurons over 28 days (Kasanga et al., 2023), we contemporaneously evaluated extracellular DA levels in the striatum and SN under baseline and depolarizing conditions at 2 time points, 7 and 28 days, following unilateral 6-OHDA lesion. We previously reported that extracellular DA levels were greatly diminished in lesioned striatum (Kasanga et al., 2024) within 7 days after 6-OHDA, replicating the severity of DA and TH loss seen upon PD diagnosis. We now extend this study to evaluate DA regulation in the SN and whether extracellular DA levels would increase against TH loss in striatum or SN ipsilateral to lesion, or contralateral to lesion as a compensatory mechanism to severe TH protein loss (Kasanga et al., 2023). To answer to this, we included a sham-operation group as a relevant control comparison. In the SN, wherein TH protein decreases at a slower rate than in striatum (Kasanga et al., 2023), we expected progressively diminished DA levels. Our results show extracellular DA levels were transiently augmented in the DA lesioned group, but only in the SN, not striatum, following depolarizing stimulation in the striatum. This study adds new insight into compensatory mechanisms that are engaged in the nigrostriatal pathway during neuronal loss, and the underappreciated role of DA signaling in the SN.

## Methods

### 2.1. Animals

Upon arrival, adult male and female Sprague Dawley (N=52; 30 F, 22 M) rats were housed in reverse light/dark cycle (lights on at 3:00 P.M., off at 3:00 A.M.) in a temperature-controlled room (22°C – 23°C). Initially, animals were pair housed in plastic cages (22 cm high, 45 cm deep, and 23 cm wide) with ad libitum food (Rodent Diet 5001; Lab Diet, Brentwood, MO, USA) and water. To ensure sufficient habituation to experimenter handling, animals were handled at least 2 times for 5-minute periods before any surgery or experimentation. After surgeries, animals were single housed to prevent housemates from chewing on in-dwelling cannulae. Rats were maintained in accordance with the guidelines of the Institutional Animal Care and Use Committee of Binghamton University and the “Guide for the Care and Use of Laboratory Animals” (Institute for Laboratory Animal Research, National Academies Press, 2011).

### 2.2. Surgical Procedures

As shown in **Fig. 1**, all rats received either a unilateral sham (0.9% NaCl + 0.1% ascorbic acid) or 6-OHDA (Sigma) lesion to the left medial forebrain bundle to deplete striatal DA (Dupre et al., 2013; Lanza et al., 2019). To do so, rats were anesthetized with inhalant isoflurane (2%-3%; Sigma) in oxygen (2.5 L/min) and placed in a stereotaxic instrument (David Kopf Instruments) with the incisor bar positioned at 5 mm below the interaural line. A 10-µL Hamilton syringe attached to a 26-gauge needle was lowered into the site (AP: −1.8 mm; ML: +2.0 mm; DV: −8.6 mm, relative to bregma). Either vehicle or 6-OHDA (3 μg/μL) was infused at a rate of 2 µL/min for a total volume of 4 µL. The needle remained in the injection site for 5 additional min before removal to allow for optimal diffusion.

**Figure 1.**
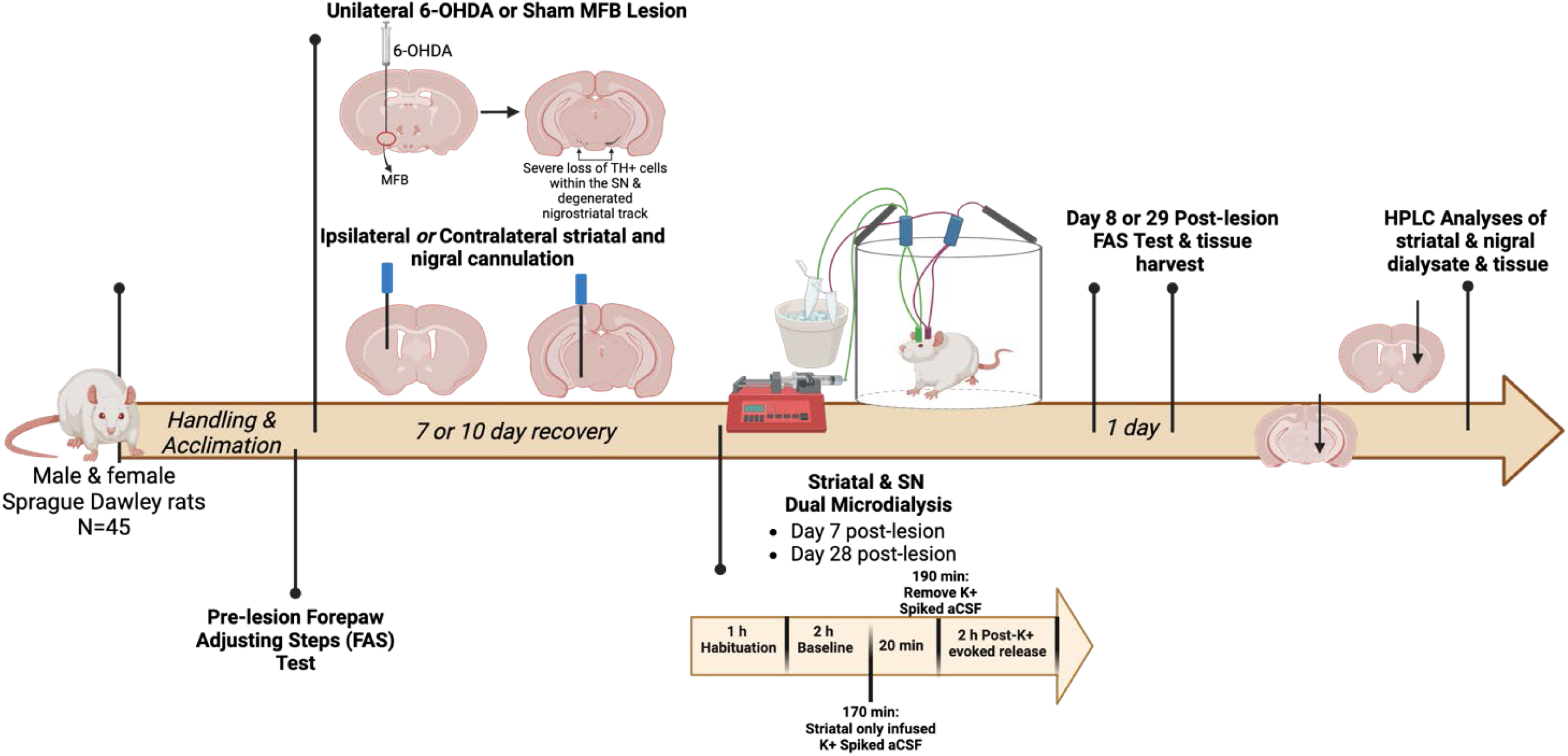
Experimental design and in vivo microdialysis timeline. Male and female Sprague Dawley rats (N=45) were acclimated to handling and the forepaw adjusting steps (FAS) test prior to surgery. During surgery, animals received either 6-hydroxydopamine (6-OHDA; n=21) or sham (vehicle; n=24) medial forebrain bundle (MFB) lesion. In the same surgery, animals were implanted with striatal and substantia nigra (SN) microdialysis cannulae either ipsilateral or contralateral to the MFB lesion or the sham operation. To assess dopamine (DA) dynamics at 7- or 28-days post-lesion, animals were either allowed to recover for 7 or 10 days. Cohorts were designated for one-time microdialysis on either day 7 or day 28 post-lesion. During testing, artificial cerebrospinal fluid (aCSF) was perfused through the striatal and nigral cannula and samples were collected every 20 min. Animals underwent 1 h of habituation and 2 h of baseline collection. With 10 min remaining of baseline (170-min timepoint) collection, striatal aCSF was replaced with a potassium chloride (KCl) spiked aCSF to evoke DA release. This infusion remained on for 20 min and replaced with aCSF again at 190 min for the remainder of testing. On day 8 or 29 post-lesion, motor impairments were assessed with the forepaw adjusting steps test (FAS) testing followed by striatal and nigral tissue collection for analysis for DA tissue content and quantification of TH protein.

During the same surgery, rats received a guide cannula (CMA 12) lowered into the striatum (ML: +/- 2.5; AP: +1.0; DV: −3.5) and substantia nigra pars compacta (ML: +/- 2.5; AP: - 5.7; DV: −7.0) and secured to the skull with surgical screws (Plastics One) and Jet Denture Repair Acrylic (Lang Dental). To assess potential contralateral effects, we counterbalanced the placement of the guide cannula with half of the rats receiving cannula placements ipsilateral to lesion and the other half receiving them contralateral to lesion. Dummy cannulae were placed in the guide to maintain patency during the recovery period. All animals that underwent surgery received one pre-operative injection of the analgesic buprenex (Buprenorphine HCl; 0.03 mg/kg, i.p; Reckitt Benckiser Pharmaceuticals Inc.) and 2 post-operative injections of carprofen (5 mg/kg, s.c.; Zoetis) 6-12 hrs apart to reduce pain and discomfort.

### 2.3. Forepaw adjusting steps test

To ascertain motor impairment in relation to the duration of lesion (or progressive nigrostriatal neuron loss as shown (Kasanga et al., 2023), we evaluated use of forelimbs. Prior to surgical intervention, all animals underwent 2 days of habituation to the forepaw adjusting steps test (FAS) followed by baseline FAS measures. Briefly, an experimenter holds both back legs and one front paw, forcing weight onto the free paw. Rats are then moved laterally across a table’s surface at a rate of 90 cm/10 seconds. In total there are 6 trials for each paw: 3 trials for the forehand movement and 3 trials for the backhand movement. At either Day 8 or Day 29 post-lesion, FAS was done to assess post-lesion motor deficits. This test is sensitive to 6-OHDA induced akinesia (Chang et al., 1999; Conti et al., 2016) and is valuable as an assay to approximate extent of striatal DA nigrostriatal lesion.

### 2.4. In vivo microdialysis

To examine the effects of nigrostriatal 6-OHDA lesion timing on extracellular DA release dynamics, rats underwent in vivo microdialysis procedures at Day 7 or 28 post-lesion (**Fig. 1**). On test days during the rats’ dark cycle (from 7:00 A.M. – 1:00 P.M.), microdialysis probes (CMA 12 Elite probe) were inserted into targeted striatal and nigral sites. Probes for the striatal site extended 3 mm beyond the cannula tip (molecular weight cut-off 100 kDA) and the nigral probes extended 1 mm beyond (molecular weight cut-off 20 kDA). In both sites, artificial cerebral spinal fluid (aCSF; 147 mM NaCl, 2.8 mM KCl (K+), 1.2 mM CaCl2, 1.2 mM MgCl2, pH 7.40 ± 0.02, sterile filtered) were perfused with a syringe pump (model 400, CMA Microdialysis) at 2 μL/min. Rats underwent 1 h habituation collection followed by a 2 h baseline collection period. At 170-min into microdialysis, potassium-chloride (K+) spiked aCSF (47.7 mM NaCl, 100 mM K+, 1.2 mM CaCl2, 1.2 mM MgCl2) was infused into the striatal site to evoke DA release. This K+-infusion remained on for 20 min, after which it was removed and replaced with aCSF again for the remainder of testing (Centner et al., 2024). Dialysate samples were taken every 20 min for 5 h and stored at −80°C until analysis. After a 24-h washout, brains were flash frozen in cold 2-methylbutane (EMD Millipore) for later tissue and protein analysis.

### 2.5 Tissue processing

Following microdialysis, striatal and nigral tissue were flash frozen for later dissection for quantification of DA tissue and TH protein levels according to established methodology (Salvatore et al., 2012a, Salvatore and Pruett, 2012). Fresh-frozen tissues were sonicated in ice-cold perchloric acid solution and centrifuged to segregate precipitated protein from the supernatant. The supernatant was analyzed in-house by HPLC (Thermofisher) for DA, dihydroxyphenylacetic acid (DOPAC) for assessment of DA turnover. A standard curve, ranging from 1.56 to 800 ng/mL was used to quantify DA and DOPAC. The resulting values for tissue DA were calculated to total recovery in the sample and normalized against total protein recovered.

The precipitated protein was sonicated in 1% sodium dodecyl sulfate solution and aliquots were assayed for total protein by the bicinchoninic acid method against an albumin standard curve. Sample buffer with dithiothreitol as a reducing agent was then added to the sonicated samples for SDS-PAGE for quantitative Western blot assessment of TH protein (Salvatore et al., 2012a). The samples for all treatments and groups were represented and balanced within each blot, so that individual differences in treatments were captured within each blot.

Tyrosine hydroxylase protein was quantified against a calibrated standard traceable over multiple studies in rats and mice (Salvatore, 2014; Salvatore et al., 2009; 2012a; 2016, 2018). Optimal total protein load for TH determination in striatum and SN was 4 and 8 µg, respectively, using the primary antibody obtained from Millipore (Temecula, CA, cat #AB152).

### 2.5 Statistics

For motor assessment, in the sham-operation group, forelimb use was evaluated using an unpaired t-test to determine whether cannulation itself affected use of the forelimb. In the 4, 6-OHDA groups (2 time points post-lesion, 2 sides (ipsilateral and contralateral to side of lesion) of cannulation) a 2-way ANOVA was used to assess if day post-lesion or side of cannulation affected percent loss of forelimb use against respective baseline use obtained pre-lesion.

For all neurochemical analyses, appropriate statistical approaches were used to determine if significant differences existed among multiple different independent variables, depending upon the comparisons made. These variables included; sham-operation, 6-OHDA lesion, impact of time of collecting microdialysate every 20 min, side of cannulation, and aCSF vs post K+-infusion conditions. In the sham-operation groups, several statistical methods were used. To analyze TH and DA tissue content in the SN and striatum, a paired t-test was used matching side of sham surgery against respective intact side within each subject. To determine impact of sham-operation on extracellular DA levels, side of cannulation, and time of microdialysate collection on extracellular DA levels, a 3-way repeated measures ANOVA was used, matching time of collection for each subject. To determine impact of K+ infusion in this group, we used a 3-way ANOVA with time after infusion, side of cannulation, and baseline vs. K+ as independent variables. Post-hoc paired t-tests were run with results obtained from the different sides of cannulation collapsed, given minimal to no impact at both day 7 (F(1,39)=3.00, *p*=0.091) and day 28 (F(1,42)=0.01, *p*=0.91) after sham-operation, to evaluate baseline vs. K+ differences at each 20 min.

As we used both male and female rats for these studies, we also determined if there were differences in extracellular DA levels in the SN and striatum in the sham-operation group using a 3-way repeated measures ANOVA, matching dialysate collection times for each rat with independent variables being days after sham surgery, sex, and time of dialysate collection. Notably, we did not find that the day after sham surgery was a significant variable in either striatum or SN, either under aCSF or following K+-infusion. We therefore collapsed the days together to evaluate if sex differences existed using repeated measures 2-way ANOVA.

For determination of lesion impact on extracellular DA, tissue DA, and total TH protein in striatum and SN, the results from each rat had to meet 2 of 3 inclusion criterion: >30% loss of forelimb use, >70% striatal DA tissue loss, and >70% striatal TH protein loss. Of 28 successfully cannulated rats that underwent 6-OHDA lesion, 7 did not meet the full criterion. These rats were excluded from statistical analysis. To ascertain impact of lesion and effect of time past lesion, a repeated measures (using time of microdialysate collection matched to each subject), a 2-way ANOVA was used including both male and female rats in the analyses as we did not find significant effect of sex in the analyses conducted in the sham-operation groups. This approach was also used to ascertain differences in percent TH and DA remaining in striatum and SN ipsilateral to 6-OHDA lesion against contralateral tissue. After verifying rats met inclusion criterion described above, the Grubb’s test was used to identify any remaining outliers with alpha set at <0.05.

## Results

### Establishment of forelimb akinesia threshold

The double cannulation procedure affected forelimb usage on the side contralateral to the cannula implant (Suppl. Fig 1). This observation compelled us to report the impact of 6-OHDA on forelimb use as a percentage of baseline (pre-lesion) noted as percent loss of forelimb use, instead of relative to forelimb use associated with the contralateral side of lesion. Average decline in forelimb use was not significantly different amongst the 4 6-OHDA-lesioned treatment groups considering (side of cannulation X day post lesion (F(1,18)=1.33, p=0.26; Suppl Fig 2)). The percent loss of forelimb use for each group was as follows (as mean ± SEM); D7 Ipsil, 45 ± 2 %; D7 Contra, 47 ± 6 %; D28 Ipsil, 48 ± 3 %; D28 Ipsil, 39 ± 4 %) (Day post lesion F(1,18) = 0.39, p=0.54; side of cannulation F(1,18) = 0.59, p=0.45). Notwithstanding the impact of double cannulation on forelimb use, sham-operation itself had no effect on forelimb use at either time point, as seen by no difference in forelimb use on the same side as the side of cannulation (Suppl. Fig. 1).

### TH expression, tissue and extracellular DA levels in SN and striatum in sham-operated rats

Sham operation had no effect on TH protein expression levels in either striatum or SN at either time point (Suppl. Fig. 3). Dopamine tissue content was not significantly affected in striatum or SN at either time point (Suppl. Fig. 4), with the exception of a modest 30% decrease in the striatum 28 days after sham operation. There were substantially greater (∼10-fold) tissue DA levels in striatum than in SN (Suppl. Fig. 4), consistent with previous studies (Kasanga et al., 2023, rev. Salvatore, 2024). Extracellular DA levels at baseline were not affected in the SN or striatum between cannulation sides (ipsil v contra (Day 7, F(1,19) = 1.47, p=0.24; Day 28, F(1,20) = 0.09, p=0.77). However, time of aCSF infusion was a factor in the SN on day 7 (time, Day 7, F(5,87) = 2.98, p=0.016; but not Day 28 (F(5,81) = 0.69, *p*=0.63; Suppl. Fig. 5A). Across all time points collected during aCSF infusion, extracellular DA levels were lower by ∼2-fold in the SN vs striatum at day 7 (F(1,19) = 8.72, *p*=0.008; Suppl. Fig. 5A) and day 28 (F(1,20) = 13.2, *p*=0.002; Suppl. Fig. 5B). This difference was reflected in the average DA levels at day 7 ((F(3,20) = 32.0, *p*<0.0001; Fig. 2A) and day 28 ((F(3,19) = 41.8, *p*<0.0001; Fig. 2B) post-sham operation. Notably, the disparity in DA levels between SN and striatum is far greater at the tissue than the extracellular level. At day 7 only, DA levels increased in the SN, ipsilateral to the sham-operation, which we attribute to a transient response to tissue injury from the cannulation of the SN.

**Figure 2.**
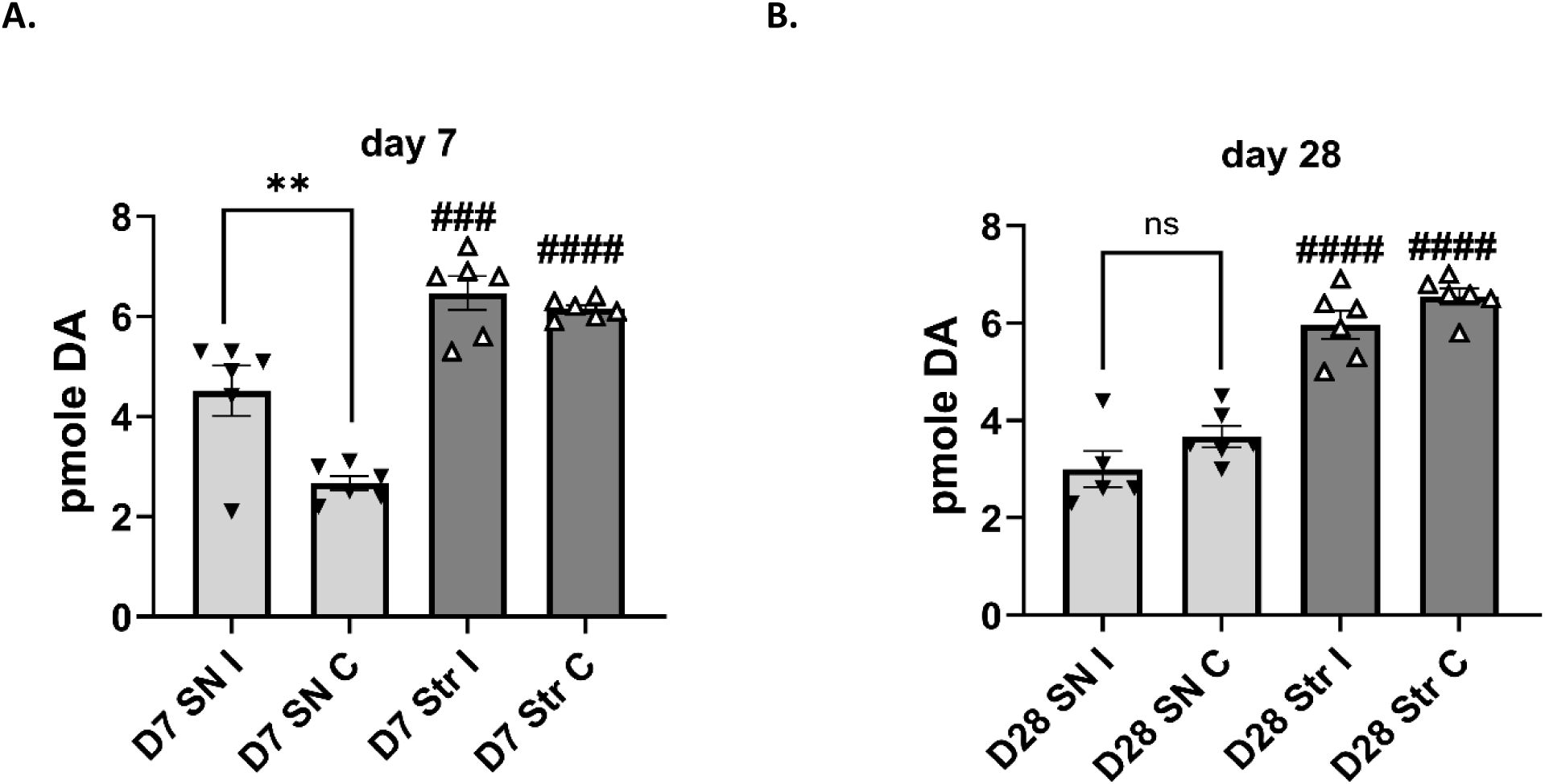
Baseline extracellular dopamine levels in substantia nigra and striatum at days 7 and 28 in sham group. **A, B.** Extracellular dopamine (DA) levels were greater in the striatum compared with substantia nigra (SN) post-sham operation (sham-op) at day 7 **(A)** and day 28 **(B). A, B.** Average extracellular DA levels, Striatum v. SN A. **Day 7.** Average extracellular DA levels were significantly different between striatum and SN in both hemispheres relative to sham-op; ipsilateral (I) (t=4.47, ^###^*p* = 0.0009, df-=20), contralateral (C) (t=7.98, ^####^*p* < 0.0001, df-=20). In the SN, there was also a significant difference in DA between the hemispheres, with extracellular DA levels ipsilateral to sham-op greater than contralateral side (t=4.24, \*\**p* = 0.002, df=20). **B. Day 28** Average extracellular DA levels were significantly different between striatum and SN in both hemispheres respective to lesion; ipsilateral (t=7.74, ^####^*p* = 0.0009, df=19), contralateral (t=7.89, ^####^*p* < 0.0001, df=19).

Following infusion of K+ into striatum, extracellular DA levels increased in striatum at both the day 7; BL v K+ (F(1,10) = 37.0, *p*<0.0001), time of infusion x BL v K+ (F(5,39)=22.1, *p*<0.0001) and time of infusion x BL v K x side of infusion (F(5,39)=2.5 *p*=0.045; Fig. 3A) and day 28; BL v K+ (F(1,10) = 34.9, p<0.0001) time of infusion x BL v K+ (F(5,42)=17.5, *p*<0.0001; Fig. 3C) and time of infusion x BL v K+ x side of infusion (F(5,42)=0.45 *p*=0.81; Fig. 3B). However at the same time in the SN, K+ infusion in striatum had no effect on extracellular DA either at day 7; BL v K+ (F(1,10) = 2.7, *p*=0.13), time of infusion x BL v K+ (F(5,43)=1.00, *p=*0.43), and time of infusion x BL v K x side of infusion (F(5,43)=2.36 *p*=0.056; Fig. 3B) or day 28; BL v K+ (F(1,10) = 1.6, *p*=0.24), time of infusion x BL v K+ (F(5,27)=1.31, *p=*0.29), and time of infusion x BL v K x side of infusion (F(5,27)=1.13, *p*=0.36) after sham-operation (Fig. 3D).

**Figure 3.**
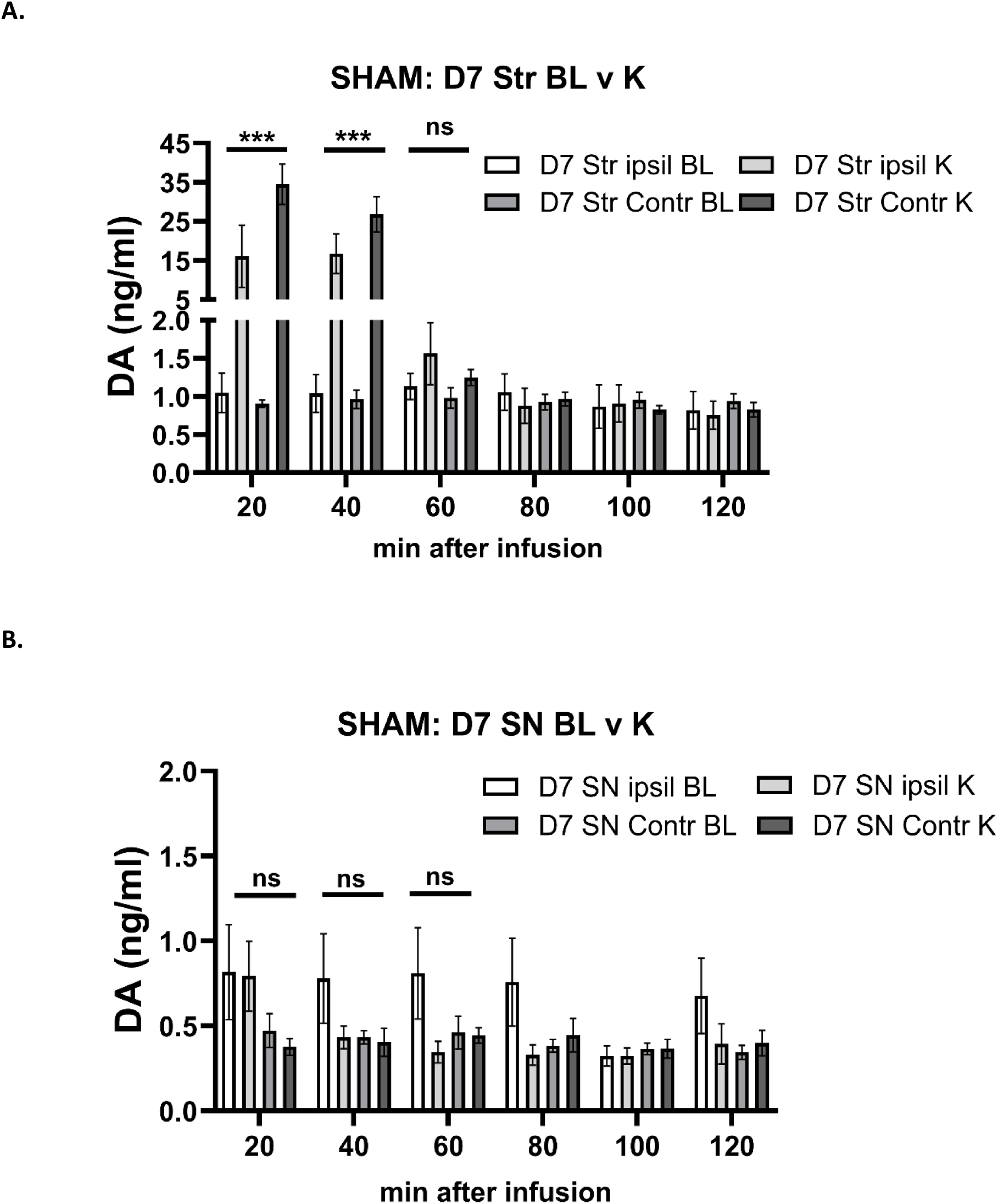
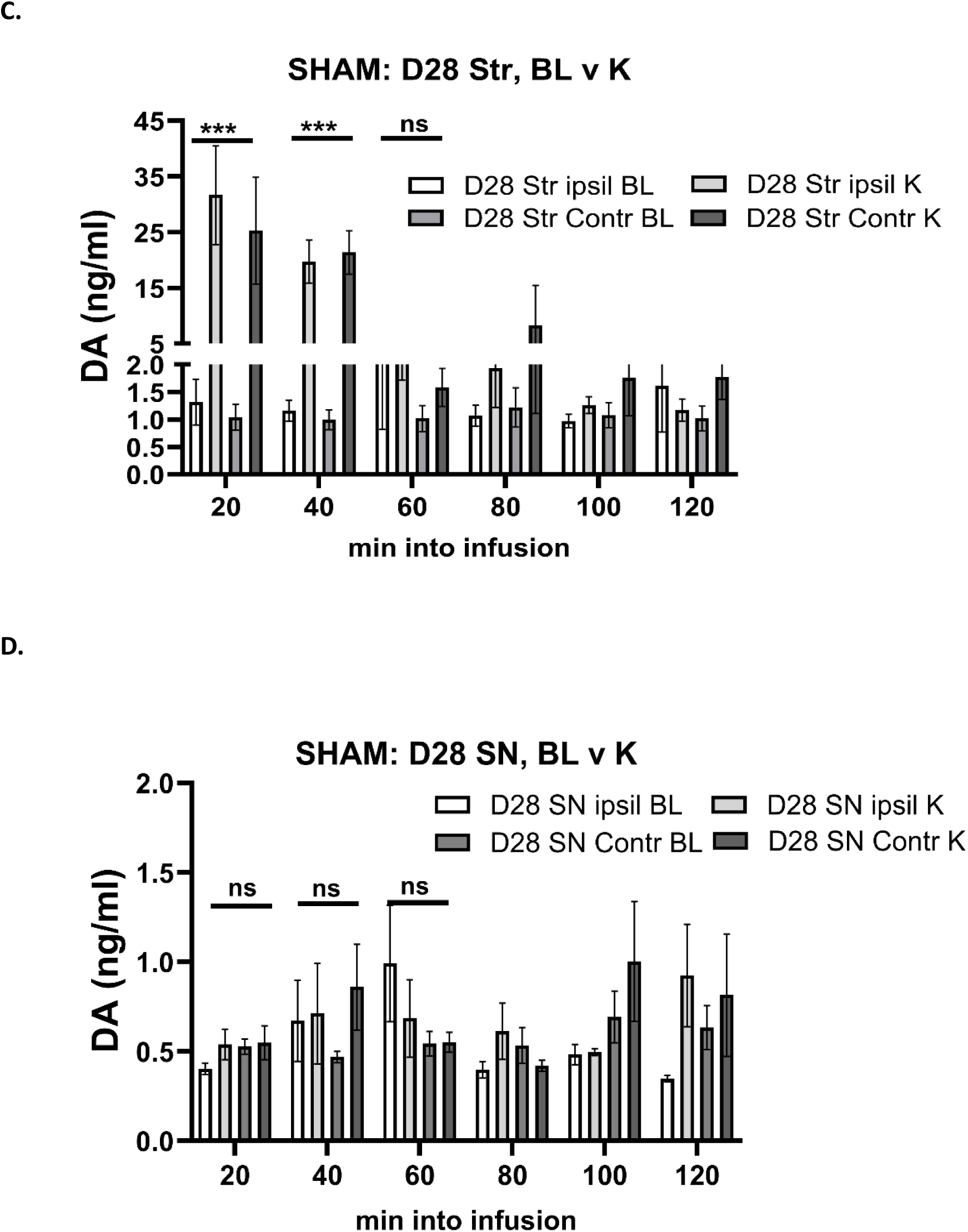
Timeline of K+-stimulated increase in extracellular dopamine levels in striatum and substantia nigra in sham group. **A, B. Day 7 after sham-operation. A. Striatum** There was a highly significant interaction between time past infusion and baseline (BL) vs K+ on extracellular dopamine (DA) levels occurring either ipsilateral or contralateral to the side of sham operation (F_(5,39)_= 22.1, *p*<0.0001), with a 15- to 35-fold increase in extracellular DA that was maintained out to at least 40 min after initiating K^+^ infusion. Significant differences in extracellular DA at 20 min (t=4.65, \*\*\**p* = 0.0002, df=20) and 40 min (t=5.28,\*\*\**p*< 0.0001, df=19), but not 60 min (t=1.49, *p* =0.15, df=19). **B. SN**. Striatal infusion of K^+^ 7 days after operation had no effect on nigral extracellular DA levels and there was no significant interaction between time of infusion and BL vs K extracellular DA levels occurring either ipsilateral or contralateral to the side of sham operation (F_(5,43)_= 1.00, *p*=0.43). **C, D. Day 28 after sham-operation. C. Striatum.** There was a highly significant interaction between time into infusion and BL vs K extracellular DA levels occurring either ipsilateral or contralateral to the side of sham operation 28 days after operation (F_(5,42)_= 17.5, *p*<0.0001), with a 15- to 25-fold increase in extracellular DA, maintained out to at least 40 min after initiating K^+^ infusion. Significant differences in extracellular DA at 20 min (t=4.81, \*\*\**p* = 0.0001, df=20) and 40 min (t=7.38, \*\*\**p*<0.0001, df=20), but not 60 min (t=1.71, *p* =0.10, df=20). **D. SN**. Striatal infusion of K^+^ 28 days after operation had no effect on nigral extracellular DA and there was no significant interaction between time of infusion and BL vs K extracellular DA levels occurring either ipsilateral or contralateral to the side of sham operation (F(5,27)= 1.13, *p*=0.29).

By 60 min after K+ infusion, the increase in extracellular DA seen in striatum returned to baseline levels (Fig. 3A, C). The side of double cannulation relative to sham-operation (ipsilateral or contralateral) had no influence on striatal DA levels (day 7, (F(1,39) = 3.0, *p*=0.091); day 28 (F(1,42) = 0.01, *p*=0.91) or nigral DA levels (day 7, (F(1,43) = 2.2, p=0.15); day 28 (F(1,27) = 0.07, p=0.79). Together, these results confirm that in the absence of nigrostriatal lesion, basal extracellular DA levels are ∼2-fold greater in striatum vs SN, and that the depolarizing influence of striatal K+-infusion to increase extracellular DA is restricted to the striatum.

### Determination of sex differences in extracellular DA under baseline and K+-infusion

Potential sex differences in extracellular DA were evaluated in the sham-operation group. Extracellular DA levels in the striatum were not affected by day after sham-operation (F=(1,19)=0.07, *p*=0.79), nor were there interactions with the other 2 variables (dialysate collection time or sex). Collapsing the results from both days after sham-operation, we did not find any significant differences due to sex (F=(1,21)=1.02, *p*=0.32), or interaction with time after dialysate collection (F=(5,91)=0.75, *p*=0.59). Given no effect of time of dialysate collection, results from each rat were averaged. No sex differences existed in striatum for extracellular DA levels (Suppl Fig. 6A). In the SN, extracellular DA levels were not affected by day after sham-operation (F=(1,20)=1.92, *p*=0.18), nor were there interactions with the other 2 variables (dialysate collection time or sex). Collapsing the results from both days after sham-operation, we did not find any significant differences due to sex (F=(1,22)=3.48, *p*=0.075), or interaction with time after dialysate collection (F=(5,98)=082, *p*=0.54). Comparison of average extracellular DA levels between sexes, we found no significant differences between sexes in the SN (Suppl Fig 6B).

Following K+-infusion, DA levels in the striatum were not affected by day after sham-operation (F=(1,19)=0.30, *p*=0.59), nor were there interactions with the other 2 variables (dialysate collection time or sex). Collapsing the results from both days after sham-operation, we did not find any significant differences due to sex (F=(1,21)=1.29, *p*=0.27), or interaction with time after K+-infusion (F=(2,39)=0.61, *p*=0.55), although there was, expectedly, an effect of time past K+-infusion (F=(2,39)=35.4, *p*<0.0001) (Suppl Fig 6C). In the SN, DA levels in the striatum were not affected by day after sham-operation (F=(1,18)=3.00, *p*=0.10), though there was an interaction between time past K+-infusion and sex (F=(2,31)=6.35, *p*=0.005). Collapsing the results from both days after sham-operation, time past K+-infusion was significant (F=2,34)=5.02, *p=*0.012), and although no significant differences were seen between sexes (F=(1,20)=2.12, *p*=0.16), there was significant interaction between time after K+-infusion and sex (F=(2,34)=6.02, *p*=0.006) due to differences isolated at day 28 post sham-operation, specifically and only at 40 min past K+-infusion (Suppl Fig 6D).

### Lesion impact on TH protein expression and DA tissue content

The 6-OHDA lesion predictably produced major TH protein loss in both striatum and SN, but with differential impact in these regions. Loss in striatum exceeded 86% by day 7, reaching 97% by day 28 (F(1,16) = 133, *p* < 0.0001), although there was no effect of time post-lesion in TH quantity in either contralateral or ipsilateral to lesioned side (F(1,16) = 1.24, *p* = 0.28; Fig. 4A). In the SN, TH loss was ∼70% by day 7 (F(1,15) = 24.9, *p* = 0.0002), with no further decrease by day 28 (F(1,19) = 0.14, *p* = 0.72; Fig. 4B).

**Figure 4.**
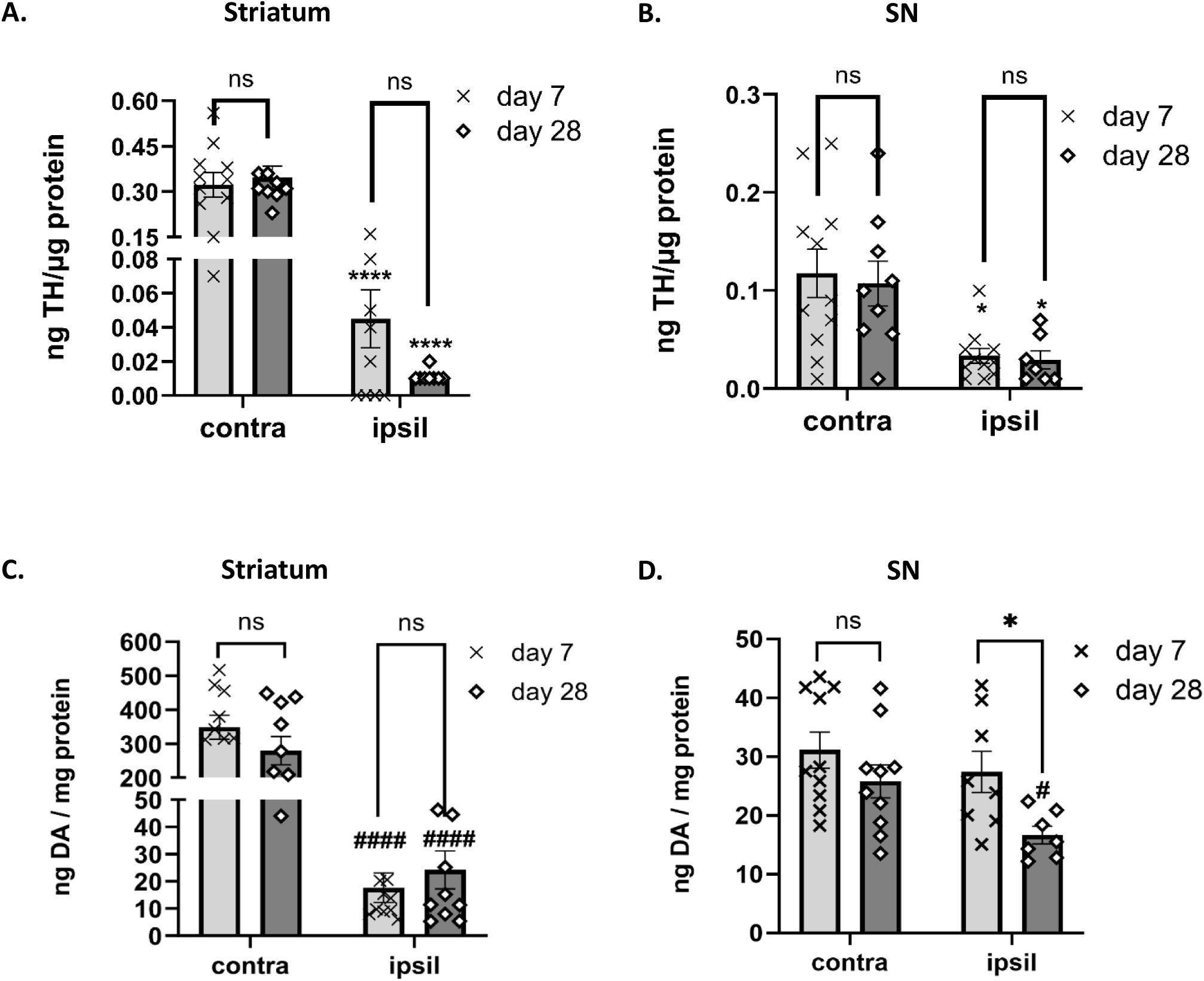
Loss of TH protein expression vs DA tissue loss in striatum vs SN during lesion progression. **A, B, TH protein loss, C,D, DA tissue loss A. Striatum TH** There was substantial TH protein loss after 6-OHDA lesion induction at 7 (t=6.0, \*\*\*\**p*< 0.0001, df=19, (86%)) and 28 days (t=8.3, \*\*\*\**p*< 0.0001, df=15 (97%). No difference in TH protein was observed between days post lesion for either the contralateral ((contra) t=0.17, *p*=0.87, df=19)) or ipsilateral ((ipsil) t=1.87, *p*=0.078, df=17) side to lesion. **B. SN TH.** There was TH protein loss after 6-OHDA lesion induction at 7 (t=2.9, \**p*= 0.011, df=19, (74%)) and 28 days (t=2.8, \**p* =0.013, df=14, (72%). No difference in TH protein was observed between days post lesion for either the contralateral ((contra) t=0.17, *p*=0.87, df=19)) or ipsilateral ((ipsil) t=1.87, *p*=0.078, df=17) side to lesion. **C. striatum DA.** There was substantial (>90%) DA loss after 6-OHDA lesion induction at 7 (t=9.2, ^####^*p*< 0.0001, df=18), and 28 days (t=6.0, ^####^*p*< 0.0001, df=19). No difference in DA was observed between days post lesion for either ipsilateral ((ipsil) t=0.75, *p*=0.46, df=18), or contralateral ((contra) t=1.26, *p*=0.22, df=18) side to lesion. **D. SN DA.** There was no DA loss after 6-OHDA lesion induction at 7 days (t=0.80, *p*= 0.43, df=16), but a 36% reduction at 28 days (t=2.53, ^#^*p* =0.023, df=15) as compared to the side contralateral to lesion. The loss of DA on the lesioned side at day 28 vs day 7 was also significantly greater (t=2.67, \**p*=0.019, df=19).

The 6-OHDA lesion also decreased DA tissue levels, but in a disparate way between striatum and SN, with more severe loss in striatum on both days after lesion. Loss in striatum exceeded 90% by day 7 and did not worsen by day 28 (F(1,18) = 116, *p* < 0.0001; Fig. 4C). However, in the SN, no DA tissue loss occurred at day 7, but by day 28, there was moderate (∼30%) DA loss (F(1,13) = 5.8, *p* = 0.031; Fig. 4D).

The magnitude of TH loss in striatum exceeded that in the SN during the course of lesion progression (F(1,30) = 9.6, *p* = 0.004; Fig. 5A). This differential loss between TH and DA was reported in our recent study (Kasanga et al., 2023) and reflects the more rapid rate of striatal TH loss compared to that in the SN in human PD and established PD models (Bezard et al., 2001; Kordower et al., 2013). The loss of tissue DA levels in SN was far less than that in striatum during the course of lesion progression (F(1,14) =56.6, *p* < 0.0001; Fig. 5B). Thus, despite significant TH protein loss by day 7 engagement of a compensatory process augmented DA signaling to diminish the impact of TH protein loss on DA levels; however, this disparate degree of DA vs. TH loss was isolated to the SN, whereas comparable levels of TH and DA loss occurred in striatum.

**Figure 5.**
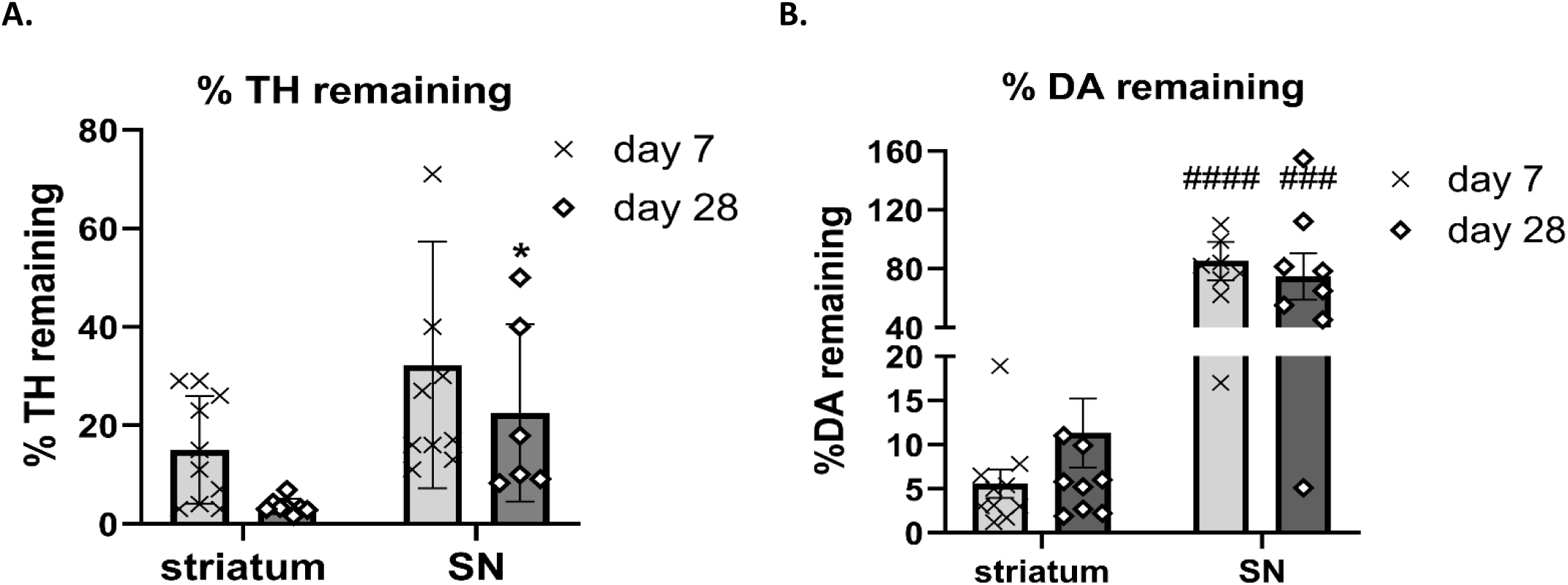
A. Tyrosine hydroxylase protein remaining striatum vs substantia nigra. Remaining tyrosine hydroxylase (TH) protein in striatum vs substantia nigra (SN) was not significantly different (t=2.0, *p*=0.061, df=18) on day 7 but was significantly different with more TH protein remaining in SN by day 28 (t=3.0, *p*=0.011, df=12). **B. Dopamine remaining striatum vs SN.** Remaining dopamine (DA) tissue levels were greatly different between striatum vs SN at both time points, with substantially more remaining in SN at day 7 ((85 v 5.6%), t=6.4, ^####^*p*<0.0001, df=17), day 28 ((74 v 11.3%), t=2.0, ^###^*p*=0.0005, df=16).

### Differential impact of lesion on baseline and K+-evoked extracellular DA

With the establishment of baseline extracellular DA levels between striatum and SN (Fig. 2) and ascertaining that K+-dependent depolarization in striatum increased extracellular DA only in striatum in the sham-operation group (Fig. 3), we determined the effect of progressive nigrostriatal neuron loss (Kasanga et al., 2023) on baseline and K+-evoked extracellular DA levels in the lesioned and contralateral to lesioned striatum and SN.

Baseline extracellular DA levels did not differ between the two time points after lesion, respective to side of lesion, in striatum (ipsil (F(1,53) = 0.00, p = 0.98); contra (F(5,39) = 0.89, p = 0.50), nor in the SN (ipsil (F(1,54) = 0.09, p = 0.75); contra (F(1,39) = 1.03, p = 0.31). However, there were significant differences in baseline extracellular DA levels between the ipsilateral and contralateral sides relative to lesion in the striatum and SN at day 7 (striatum, (F(2,69) = 11.8, p<0.0001; Suppl Fig. 7A); SN, (F(2,67) = 3.9, p =0.026; Suppl Fig. 6B) and in only in striatum at day 28 post lesion (striatum, (F(2,69) = 17.2, p <0.0001; Suppl Fig. 6C); SN, (F(2,68) = 0.96, p =0.39; Suppl Fig. 6D). Specifically, in striatum, on day 7 after 6-OHDA lesion, extracellular DA levels in the side contralateral to lesion were increased compared to the sham-operated group (q=3.98, p=0.017) and to a greater extent compared to striatum ipsilateral to lesion (q=6.75, *p*<0.0001; Suppl Fig 6A). Given the differences in baseline levels in striatum between contralateral to lesion v sham-operated group, we evaluated the first 60 min following 20 min of K+infusion across both time periods after lesion or sham-operation. There was a substantial effect of time after infusion (F (2,18)=50.8, *p*<0.0001), but time after lesion or sham-operation was not a significant factor (F (1,9)=1.0, *p=*0.34), nor were there differences between extracellular DA levels contralateral to lesion v sham-operation group, although there was a notable trend toward an increase in striatum contralateral to lesion (F (1,9)=4.1, *p*=0.073).

Following K+ infusion 7 days after nigrostriatal lesion, extracellular DA levels increased in the striatum contralateral to lesion (F(1,8) = 61.2, p < 0.0001), time of infusion x lesion (F(11,83) = 25.0, p<0.0001; Fig. 6A). Extracellular DA was 30-fold greater than baseline conditions in the striatum contralateral to lesion within the first 60 min after K+ infusion (F(1,22) = 52.1, p < 0.0001; Fig. 6B). In contrast, on the lesioned side, differences in extracellular DA versus baseline after K+ were diminished > 90%, with only a modest 2-fold difference above baseline within the first 60 min after K+-infusion (F(1,22) = 8.6, p = 0.012; Fig. 6C), and no significant difference at any 20 min interval. These disparities were similar to the stark loss in DA tissue levels resulting from lesion. These results indicate that a compensatory response to augment extracellular DA levels does not exist to counteract severe DA tissue loss, and that tissue DA levels exert control over extracellular levels.

**Figure 6.**
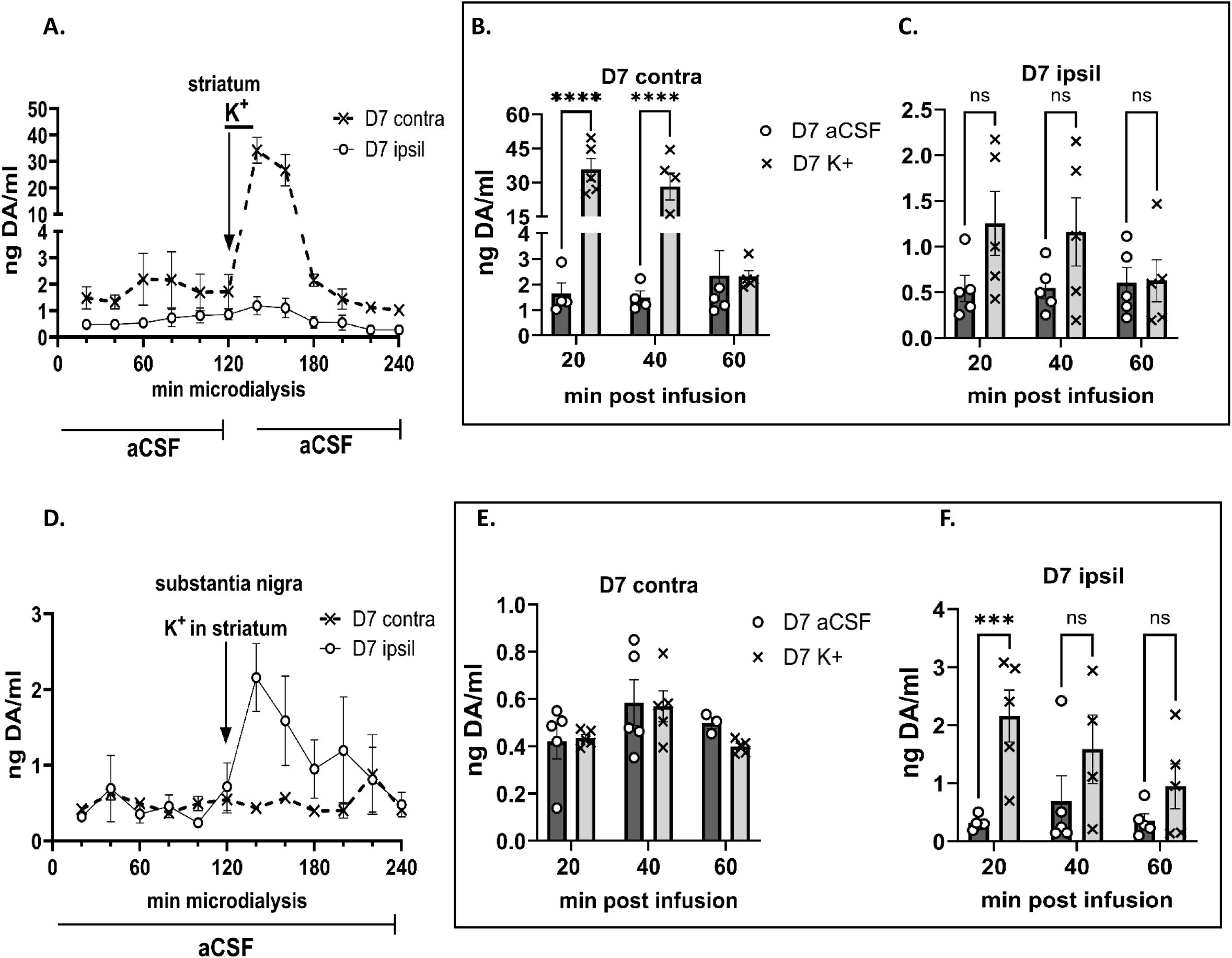
Depolarization-stimulated dopamine levels, 7 days post-lesion, aCSF v K+ matched comparison. **. Striatum (A, B, C), Substantia nigra (D, E, F). Striatum A.** K+-evoked extracellular dopamine (DA) levels were intact in the striatum, contralateral to lesion, increasing ∼30-fold within the first 20 min after K+-infusion. However, in striatum ipsilateral to lesion, K+-evoked extracellular DA levels were virtually eliminated (Ipsil vs contra, (F(1,8)=61.2, *p*<0.0001)). **B. Contralateral to lesion.** 20 min (t=6.87, \*\*\*\**p*<0.0001); 40 min (t=5.39, \*\*\*\**p*<0.0001); 60 min (t=0.01, *p*=0.99). **C. Ipsilateral to lesion.** 20 min (t=2.68, *p*=0.059); 40 min (t=2.33, *p*=0.115); 60 min (t=0.08, *p*=0.99). **SN. D.** K+-evoked extracellular DA levels increased following striatal K+-infusion were restricted to the SN, ipsilateral to lesion (Ispil vs contra (F(1,9)=5.56, *p=*0.043)). **E. Contralateral to lesion.** No significant differences in extracellular DA were observed at any time point after striatal K+-infusion. **F. Ipsilateral to lesion.** 20 min (t=5.25, *p*=0.0008); 40 min (t=2.28, *p*=0.132); 60 min (t=1.70, *p*=0.35).

In the SN, there were significant differences between the lesioned and contralateral to lesion SN extracellular DA following striatal K+ infusion (F(1,9)=5.6, *p* = 0.043; Fig. 6D). However, the differences were opposite to those seen contemporaneously in the striatum. Similar to the lack of effect of K+ infusion in the SN of the sham-operation group, extracellular DA levels in the SN contralateral to lesion were not different between baseline and after striatal K+ infusion (F(1,8) = 2.41, *p* =0.157; Fig. 6E). However, in the SN ipsilateral to lesion, there were highly significant increases in extracellular DA in the SN after K+-infusion in striatum (F(1,11) = 27.9, p = 0.0003; Fig. 6F), with extracellular DA increasing by ∼7-fold within the first 20 min.

By 28 days post-lesion, following K+ striatal infusion, extracellular DA levels still increased substantially in the striatum, contralateral to lesion compared to lesioned side (F(1,8) = 82.9, *p* < 0.0001; Fig. 7A). Notably, extracellular DA was still 30-fold greater than baseline conditions, contralateral to lesion, within the first 60 min after K+ infusion (F(1,8) = 91.6, *p* < 0.0001; Fig. 7B). In contrast, in the ipsilateral lesioned side, differences in extracellular DA versus baseline after K+ were still diminished by > 90%, with no significant difference in levels between BL and following K+ infusion (F(1,25) = 3.77, *p* = 0.064; Fig. 7C).

**Figure 7.**
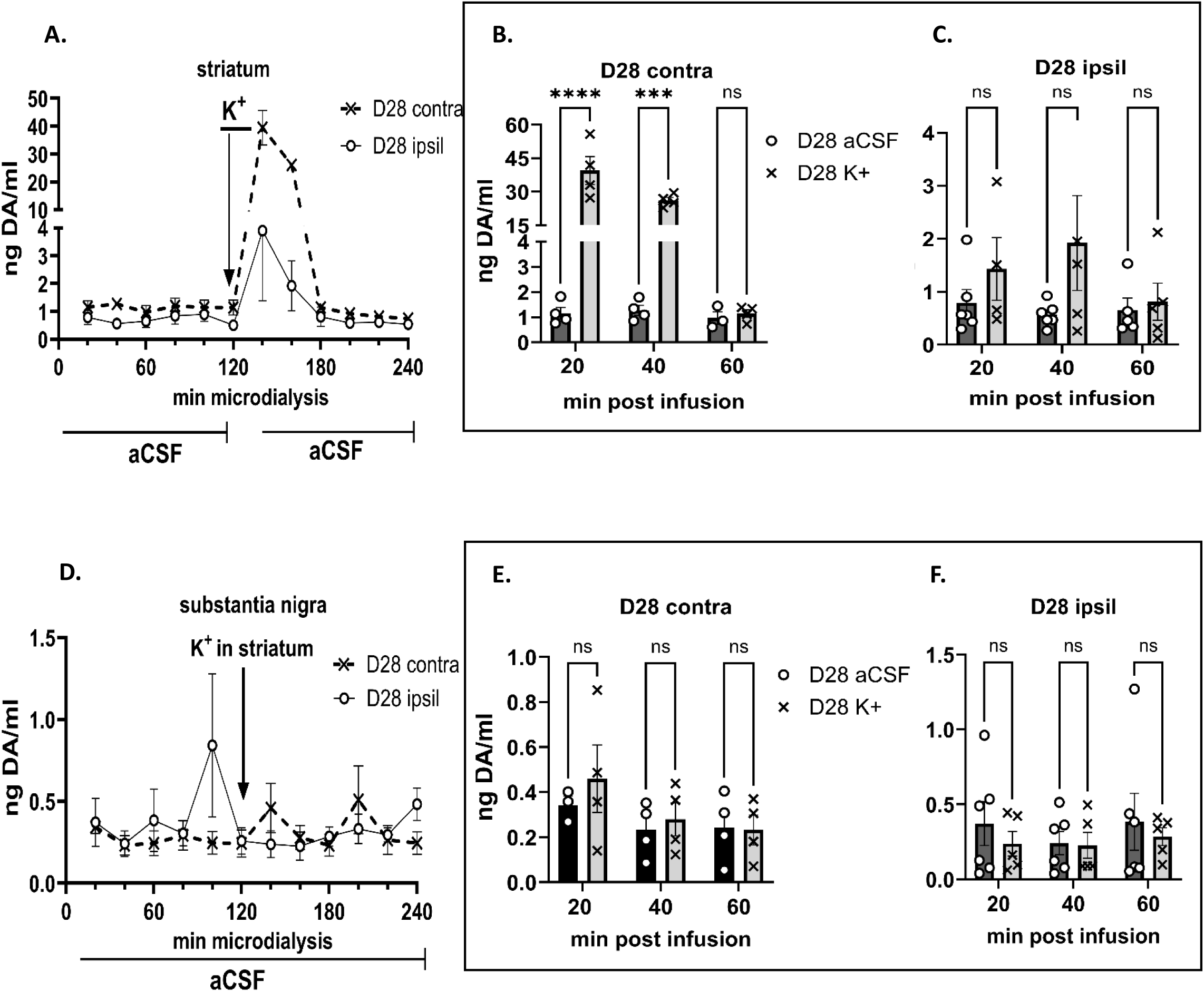
Depolarization-stimulated dopamine levels, 28 days post-lesion, aCSF v K+ matched comparison. **Striatum (A, B, C), Substantia nigra (D, E, F). Striatum A.** K+-evoked extracellular dopamine (DA) levels were intact in the striatum, contralateral to lesion, increasing ∼30-fold within the first 20 min after K+-infusion. However, in striatum ipsilateral to lesion, K+-evoked extracellular DA levels were virtually eliminated. Ipsil vs contra (F(1,8)=82.9, *p*<0.0001). **B. Contralateral to lesion.** 20 min (t=10.3, \*\*\*\**p*<0.0001); 40 min (t=6.69, \*\*\*\**p*=0.0005); 60 min (t=0.01, *p*=0.99). **C. Ipsilateral to lesion.** 20 min (t=0.97, *p*=0.99); 40 min (t=2.19, *p*=0.113); 60 min (t=0.04, *p*=0.99). **Substantia nigra. D.** K+-evoked extracellular DA levels did not change following striatal K+-infusion in either side relative to lesion (F(1,9)=0.20, *p=*0.67). **E. Contralateral to lesion.** No significant differences in extracellular DA were observed at any time point after striatal K+-infusion. **F. Ipsilateral to lesion.** No significant differences in extracellular DA were observed at any time point after striatal K+-infusion.

In the SN, by day 28 post-lesion, the significant differences between the lesioned and contralateral to lesion following striatal K+ infusion seen at day 7 were no longer apparent (F(1,8) = 0.20, p= 0.67; Fig. 7D). Extracellular DA levels in the SN contralateral to lesion were not significantly different between baseline and after striatal K+ infusion (F(1,8) = 1.64, p=0.24; Fig. 7E). On the lesioned side, unlike at day 7, there was no longer a difference between extracellular DA levels at baseline vs that following K+ infusion (F(1,12) = 0.94, p=0.35; Fig. 7F).

To compare across the two time points after lesion, responses to K+ in striatum and SN ipsilateral to lesion were compared on day 7 vs. day 28. There were no differences in extracellular DA levels within the first 60 min after striatal K+ infusion between the day 7 and day 28 cohorts (F(1,25) = 0.66, p=0.42; Fig. 8A), suggesting there was little change in the major decrease in extracellular DA during nigrostriatal lesion progression. However, in the SN, the difference in depolarization-stimulated DA levels as lesion progressed was substantial, with significantly greater DA levels after striatal K+ infusion early (day 7) versus late after lesion (day 28) (F(1,25) = 22.0, p<0.0001; Fig. 8B).

**Figure 8.**
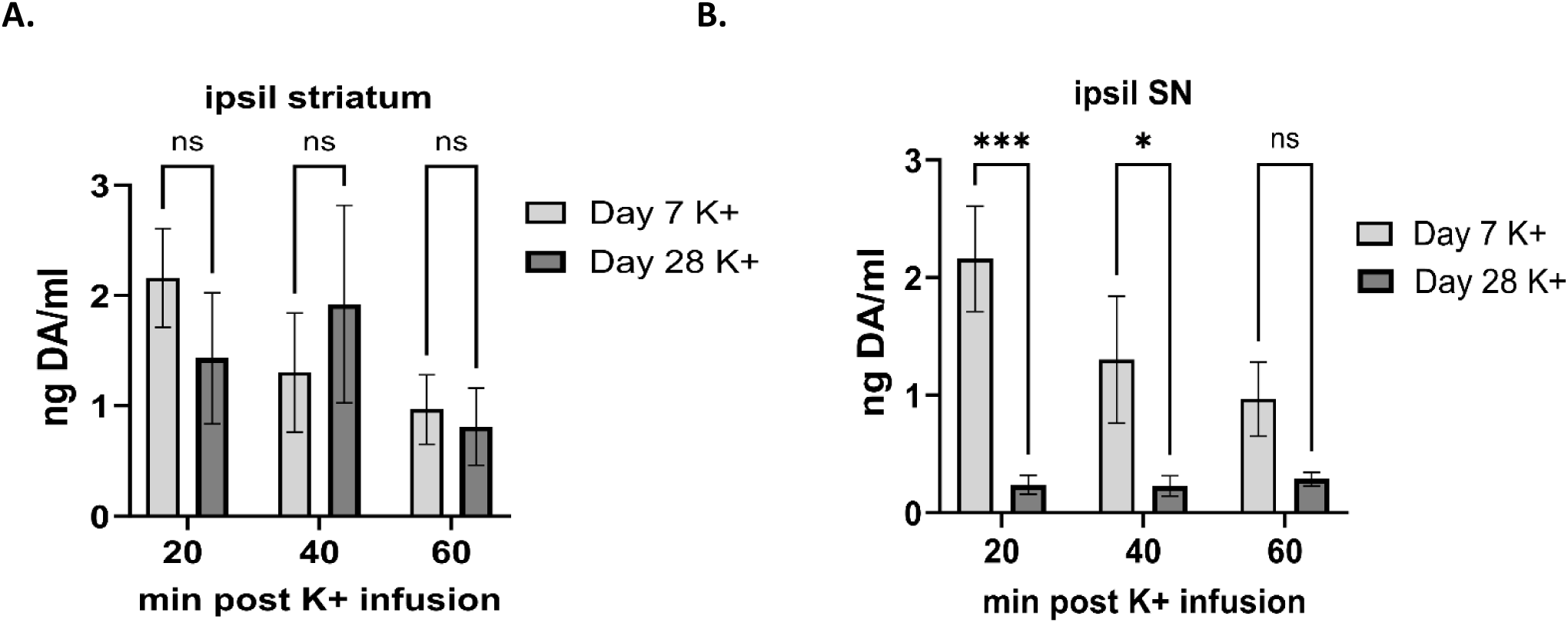
Compensatory increase in extracellular dopamine is transient and restricted to the substantia nigra. **A. Striatum** There was no significant difference in extracellular dopamine (DA) in striatum after depolarizing stimulation therein as a function of time past lesion induction. **B. Substantia nigra.** There is a transient increase in extracellular DA in the substantia nigra (SN) after depolarizing stimulation, as a function of time past lesion induction. 20 min (t=4.19, *p*=0.0003); 40 min (t=2.34, *p*=0.027); 60 min (t=1.55, *p*=0.13).

## Discussion

It has been well-established that the motor impairments driven by nigrostriatal neuron loss in PD do not occur until there is ∼50% loss of these neurons, which is associated with 80% loss of TH protein in the striatum (Bezard et al., 2001; Johnson et al., 2018; Kordower et al., 2013; Tabbal et al., 2012). Compensatory mechanisms have been thought to be engaged to augment DA signaling in the striatum against neuron or TH protein loss (Blesa et al., 2012; 2017; Pifl and Hornykiewicz, 2006); this includes augmentation of DA uptake through catecholamine transporters other than DA transport (Chotibut et al., 2012, 2014a). Our results now show that extracellular DA levels are not augmented when TH protein loss in striatum exceeds 80%, even under depolarizing conditions. Notably, loss of extracellular DA under depolarizing conditions was commensurate with drastic loss of tissue DA and TH protein, indicating that TH protein itself plays a significant role in any augmentation of DA signaling in striatum. In contrast, both extracellular and tissue DA levels in the SN did not coincide with significant TH protein loss therein, suggesting that both post-translational modification of TH and release capacity were augmented to diminish the impact of TH protein loss on DA signaling in the SN. Perhaps most strikingly, this increase in extracellular DA in the SN following depolarizing stimuli occurred only during the earlier stages of progressive nigrostriatal neuron loss, as previously shown in this model (Kasanga et al., 2023).This indicates that compensatory mechanisms to augment DA signaling in the nigrostriatal pathway during progressive neuronal loss may occur at the extracellular level only in the SN and not striatum, even though the origin of stimulation occurred in striatum (Salvatore, 2024).

Our results show that in addition to SN-restricted increased DA biosynthesis during nigrostriatal neuron loss (Kasanga et al., 2023), plasticity in DA signaling therein extends to regulation at the extracellular level in SN, with a transient increase in synaptic levels of DA. This increase only arises from depolarizing stimulating in the lesioned striatum, suggesting that the trigger to increase DA levels in the SN results from alterations in basal ganglia circuitry that begin in the striatum resulting from major DA loss therein. Augmentation of DA signaling is thought to prevent onset of parkinsonian signs until there is at least 70-80% loss of TH or dopamine transporter (DAT) in the striatum (Blesa et al., 2017; Furukawa et al., 2022; Pifl and Hornykiewicz, 2005; Zigmond, 1997). However in the striatum, DA tissue levels decrease to the same magnitude as TH protein (Kasanga et al., 2023), suggesting no augmentation of DA biosynthesis therein when TH loss is at the magnitude seen at PD diagnosis (Kordower et al., 2013; Nandhagopal et al., 2011). In the current study, we found essentially the same outcome, comparable TH protein and DA tissue loss now further showing that extracellular DA does not increase despite TH and DA tissue loss in striatum. Notably, it is plausible that by our study design we may have missed augmented extracellular DA levels that might have occurred prior to 7 days after lesion induction. That said, our data indicate that possible enhancement of DA release does not occur when TH protein loss meets the threshold of loss that occurs at PD diagnosis. Whereas there were robust increases in DA in the striatum under depolarizing conditions in the sham group and contralateral to lesion in the 6-OHDA group, there was a decrease in extracellular DA at the baseline which was greatly magnified under depolarizing conditions; in fact DA levels were nearly no different (∼1.5-fold) than levels at baseline after K+-infusion. We also did not find a significant difference in extracellular DA after depolarizing stimulation in the striatum contralateral to lesion compared to sham-operation group at either time point, indicating that the slight increase in baseline levels did not augment potential DA release capacity therein. This new finding suggests that if release capacity was augmented, as previously reported (Perez et al., 2008), it appears that the inherent DA tissue content in striatum ultimately dictates extracellular levels, including release capacity. Thus, even if DA release mechanisms were autonomously affected from biosynthesis or vesicular packaging, the increase would be minimal, potentially far below the levels needed for normal motor function.

The source of elevated levels of extracellular DA may be explained by changes in basal ganglia circuitry induced by the loss of nigrostriatal DA (Blesa et al., 2017; 2022). The neuronal pathways that comprise the indirect pathway include the final neuronal pathway which targets the SN originating from the subthalamic nucleus (STN); the subthalamonigral pathway. This glutamatergic pathway innervates the GABAergic output neurons from the SN pars reticulata (SNr) and becomes overactive following the loss of DA in striatum. As a result, there is an increase in the tonic inhibitory output of the basal ganglia by the STN, which may also overexcite the remaining nigrostriatal neurons, leading to increased DA release. Several lines of evidence indicate that this pathway does innervate the nigral DA neurons (Iribe et al., 1999; Lee and Tepper, 2009), and extracellular glutamate levels increase in the SN in a mouse PD model (Meredith et al., 2009). A glutamatergic basis for augmented tissue and extracellular DA in the SN may be initiated from the influence of depolarizing stimulation arising from the subthalamic nucleus (Mintz et al., 1986; Iribe et al., 1999). In response to depolarizing stimulation or modulation of glutamate receptor signaling or uptake, Ser19 TH phosphorylation levels are changed (Lindgren et al., 2000; Chotibut et al., 2014b; Salvatore et al., 2012b, 2014; Salvatore and Pruett, 2012). Consistent with our findings here, ser19 TH phosphorylation levels increase within a week after the induction of 6-OHDA lesion (Salvatore, 2014). In summary, the source of increased extracellular DA and tissue DA, despite TH protein loss, may arise from overactive glutamate release from the subthalamic nucleus. Furthermore, while it is unlikely storage capacity is increased in the striatum (Nandhagopal et al., 2011), DA vesicular storage may slightly increase in the SN for somatodendritic release early post-lesion, resulting in the increase of release during K+ striatal infusion seen in the current work. Future assessments of this circuit change could incorporate an analyses of vesicular monoamine transport 2 (VMAT2) levels, which could indicate if an upregulation of storage is occurring in addition to DA synthesis and release (Tokuoka et al., 2011).

Lastly, cross-hemispheric collaterals are an interesting component in disease models. In the present data we observed significant increases in contralateral striatal DA at baseline and following depolarizing stimulation, suggesting an intact contralateral side untouched by our MFB lesion. Work assessing cross-hemispheric projections from the SN to the striatum in a 6-OHDA lesioned model suggests a portion of these contralateral projections that survive may protect against certain medication related side effects such as L-DOPA-induced dyskinesia (Iyer et al., 2022). Knowing this, targeting these surviving neurons and respective projections may provide additional compensation for motor dysfunction.

### Limitations

In the sham-operation group, depolarizing stimulation was associated with an increase in extracellular DA that was isolated at 28, but not 7, days after sham-operation. Even more unexpected was that this increase occurred only 40 min after striatal K+-infusion. We speculate that this may be a delayed response to tissue injury, but it is interesting that this increase was seen only in females. While not the subject of our study, this preliminary finding may be a piece of evidence of why females are at less risk for PD, with inherent resiliency against CNS insults not necessarily specific to nigrostriatal neurons. This is the first study to our knowledge that has 3 coincident measures of DA signaling within the same tissues, making it possible to determine whether there is consistency in the magnitude of loss of TH protein, DA tissue content and extracellular DA in striatum and SN. However, given the nature of the experimental design to ascertain extracellular striatal and SN DA levels under both baseline and K+ infusion, we were unable to collect tissue at the time of K+ striatal infusion. Thus, it remains unknown whether tissue DA levels would be further augmented along with extracellular DA under depolarizing conditions. It is established that depolarizing stimuli can augment DA biosynthesis (Salvatore et al., 2001). Notably, the quantitation of TH protein in our study matches previously reported levels (Pruett and Salvatore, 2012; Kasanga et al., 2023, Salvatore, 2024), adding an additional degree of certainty that changes in extracellular DA levels would also be consistent across *in vivo* studies. Despite that we did not observe any increase in extracellular DA in striatum after lesion induction by day 7, it is plausible that extracellular DA levels may be less influenced by DA tissue or TH protein loss occurring earlier during neuron loss. It would be important to determine if there is a breaking point of whether the magnitude of TH protein loss would not be commensurate with DA release potential, which could identify a compensation mechanism in striatum at least in the very early stages of neuronal loss.

### Summary

Our results support more recent work that indicates pre-symptomatic compensation mechanisms that respond to nigrostriatal neuron loss, as seen in PD, do not involve increased DA signaling in the striatum (Bezard et al., 2003; Blesa et al., 2017, 2022; Kasanga et al., 2023). With the caveat that DA compensation may have occurred much earlier in striatum after lesion induction in our model, our study adds new insights that compensatory mechanisms that increase DA signaling in response to neuronal loss involve increased extracellular DA levels in the SN, and not striatum. Notably this increase is seen early, but not late, after lesion induction and occurs only with confirmed loss of TH protein. This is compelling evidence that there is a dopaminergic component in the nigrostriatal pathway to compensate for TH protein loss during neuronal loss. A strength of our study is the contemporaneous collection of extracellular DA from both striatum and the SN, which for one, drastically reduces the likelihood of study to study variation, and second, represents changes in DA signaling the entire nigrostriatal pathway during neuronal loss. Our findings align with confirmation of loss of motor function and TH protein in both striatum and SN at the magnitude seen at PD diagnosis. Given our understanding of the how basal ganglia function changes during DA loss in PD, the increase in extracellular DA in the SN may be driven by enhanced activity from the subthalamic nucleus. This possibility is supported by studies that show innervation of the DA neurons from this region and that glutamate levels increase in the SN during DA loss. Our results provide a new and important piece of the puzzle to optimize therapeutic strategies; in this case, the therapeutic potential for targeting regions outside the depleted striatum. It is conceivable that therapeutic targeting of DA release in the SN could potentially provide longer compensation to preserve motor function or delay motor impairment.

## Declarations

### Ethics approval

All animals were used in compliance with the Office of Laboratory Animal Welfare guidelines and protocols approved by the Institutional Animal Care and Use Committees and at the University of North Texas Health Science Center, Binghamton University, and the Animal Care and Use Review Office, US Army Medical Research and Development Command

### Consent for publication

Not applicable

### Availability of data and materials

Data generated or analyzed during this study are included in this published article and its supplementary information files. Additional data and materials are available from the corresponding author on reasonable request.

### Competing interests

The authors declare that they have no competing interests relevant to the content of this article, no financial or proprietary interests in any material discussed in this article, or affiliations with or involvement in any organization or entity with any financial interest or non-financial interest in the subject matter or materials discussed in this manuscript.

### Funding

This study was funded by the US Department of Defense U.S. Army Medical Research and Material Command Congressionally Directed Medical Research Program (Award Number W81XWH-19-1-0757). The funders had no role in the study design, in the collection, analysis, and interpretation of data, or writing of the manuscript.

### Authors’ contributions

Study design; MFS, CB. Study conduct; AG, RM, WN. Data collection and analysis; MFS, RM, WM, AG, SC. Interpretation of results; MFS, CB, AG. Writing manuscript; AG, MFS, EAK. Editing manuscript; MFS, CB, AG, RM. All authors read and approved the final manuscript.

## Notes

### Competing Interest Statement

The authors have declared no competing interest.

