## Supplemental Figures for "Preservation of dopamine neurotransmission during nigrostriatal neuron loss in rat Parkinson’s model: evidence for increased dopamine signaling in substantia nigra"

**Supplemental Results**

**Supp. Figure. 1**

**A.**

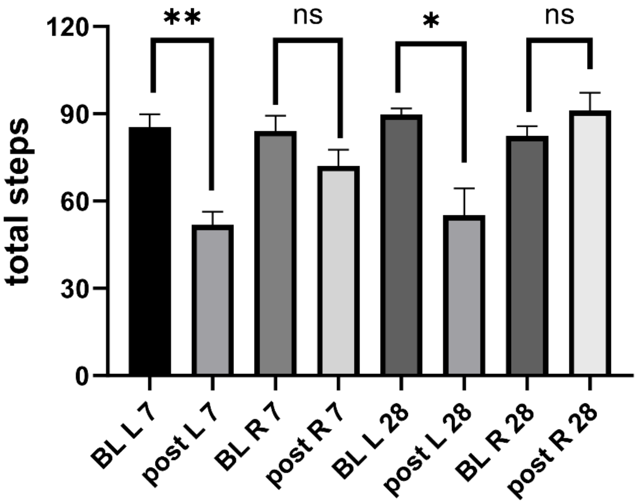


**B.**

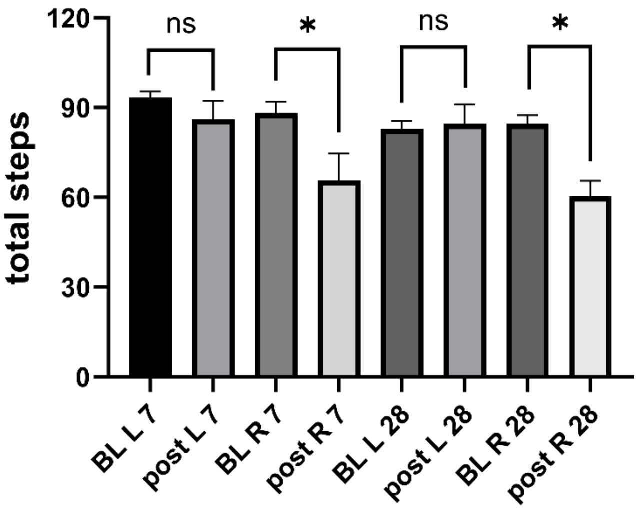


**Suppl Fig 1. Impact of contemporaneous cannulation of striatum and SN on forelimb use.** Rats of the Sham-operation group showed decreased use of the forelimb contralateral to the side of double cannulation. **A. Right side cannulation.** Use of left forelimb decreased 40% either 7 or 28 days after the sham-operation conducted on the left mfb. Left forelimb use day 7 after sham-op (post L 7) vs pre-operation baseline (BL L 7) (t=5.23, ***p*=0.0002, df=12). Right forelimb use day 7 after sham-op (post R 7) vs pre-operation baseline (BL R 7) (t=1.54, *p*=0.15, df=12). Left forelimb use day 28 after sham-op (post L 28) vs pre-operation baseline (BL L 28) (t=3.61, **p*=0.005, df=10). Right forelimb use day 28 after sham-op (post R 28) vs pre-operation baseline (BL R 28) (t=1.22, *p*=0.25, df=10). **B. Left side cannulation.** Use of right forelimb decreased 30 - 40% 7 or 28 days after the sham-operation conducted on the left mfb. Right forelimb use day 7 after sham-op (post R 7) vs pre-operation baseline (BL R 7) (t=2.32, **p*=0.039, df=7). Left forelimb use day 7 after sham-op (post L 7) vs pre-operation baseline (BL L 7) (t=1.12, *p*=0.29, df=8). Right forelimb use day 28 after sham-op (post R 28) vs pre-operation baseline (BL R 28) (t=4.03, **p*=0.002, df=10). Left forelimb use day 28 after sham-op (post L 28) vs pre-operation baseline (BL L 28) (t=0.23, *p*=0.82, df=10).

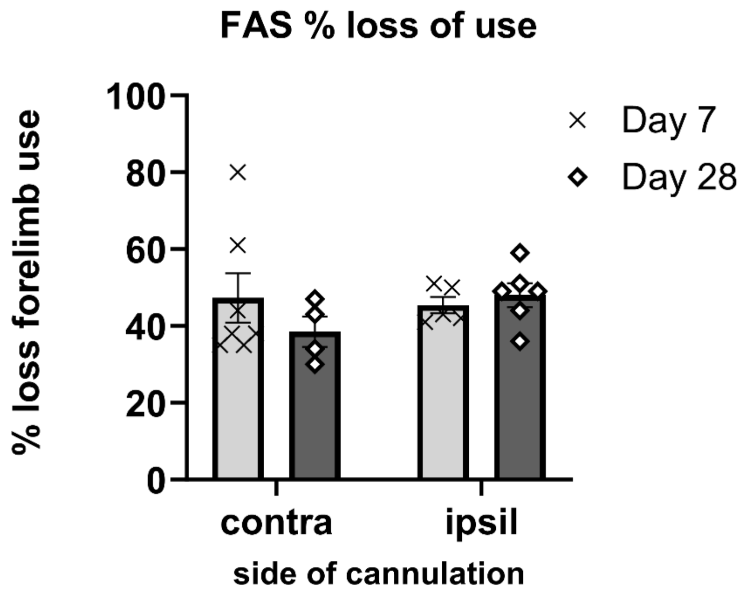


**Suppl Fig 2. Establishment of forelimb use deficits following 6-OHDA lesion induction.**

Although the double cannulation procedure decreased forelimb use on the side contralateral to cannulation (see Suppl. Fig. 1), forelimb use decreased significantly from baseline (pre-lesion) contralateral to 6-OHDA lesion side, with no significant difference as a function of time period after lesion induction Day 7 vs 28, and regardless of double cannulation side. These results establish the level of forelimb impairment associated with loss of TH protein, and changes in DA tissue and extracellular levels.

.

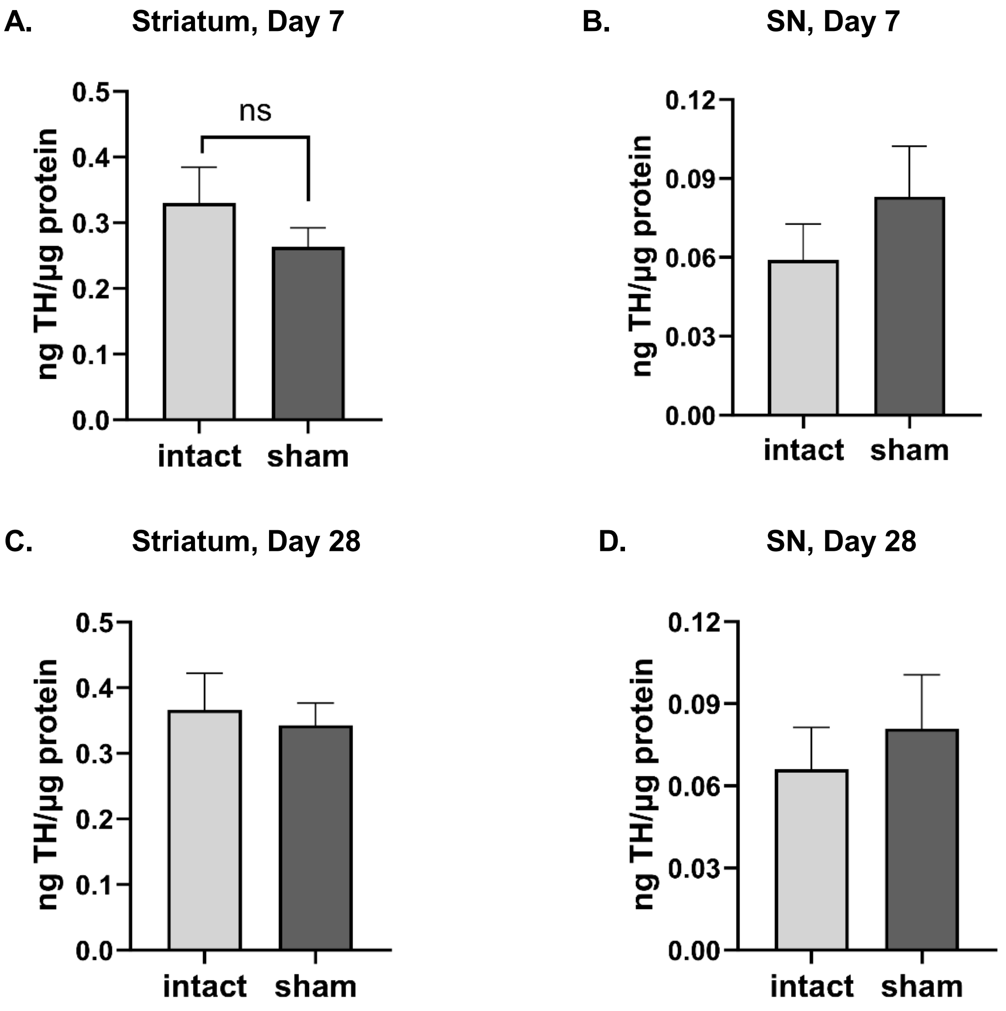


**Suppl Figure 3. TH protein expression in sham-operated groups. A, B, day 7, C,D, day 28. A. Striatum.** There was no effect on TH protein expression compared to intact striatum 7 days after sham-operation (sham) ($t=1.48$, $p=0.17$, $df=10$). **B. SN.** There was no effect on TH protein expression compared to intact SN 7 days after sham-operation (sham) ($t=1.23$, $p=0.25$, $df=9$). **C. Striatum.** There was no effect on TH protein expression compared to intact striatum 28 days after sham-operation (sham) ($t=1.77$, $p=0.12$, $df=7$). **D. SN.** There was no effect on TH protein expression compared to intact striatum 7 days after sham-operation (sham) ($t=1.42$, $p=0.19$, $df=9$).

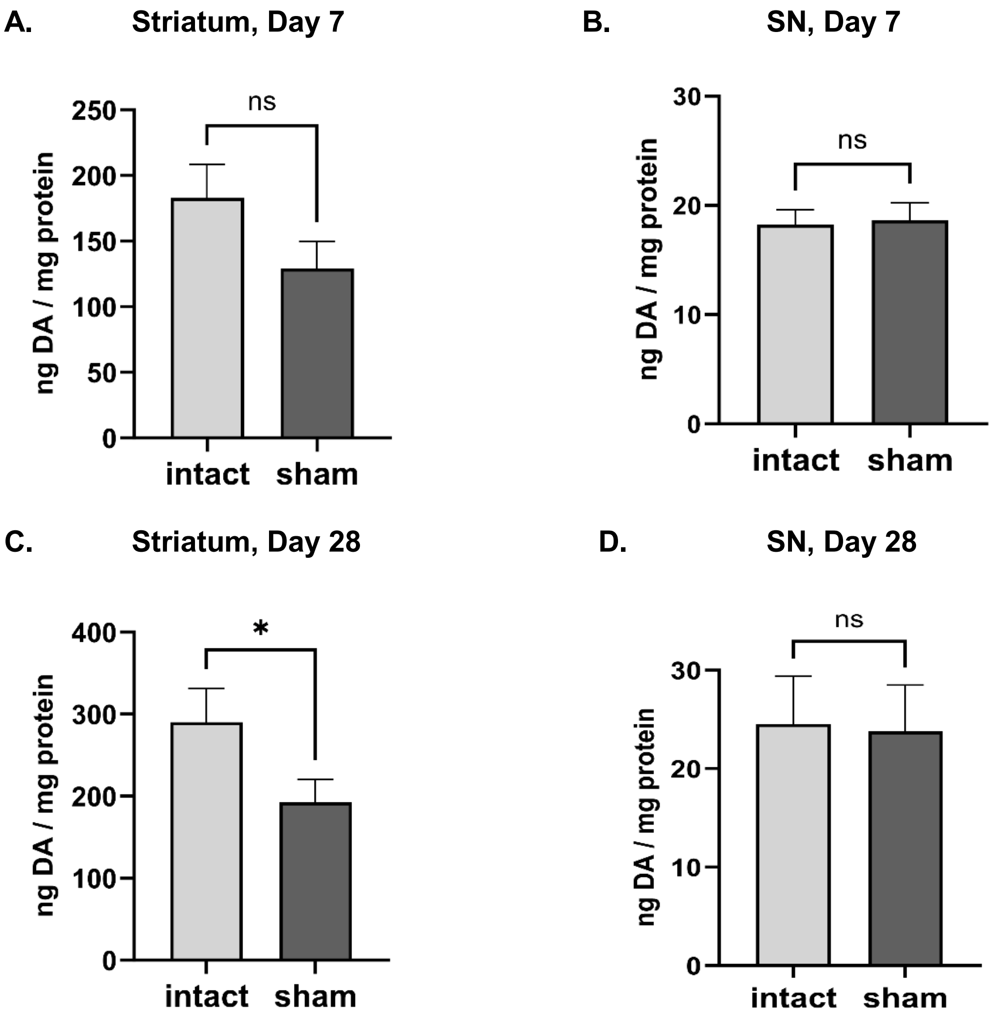


**Suppl Figure 4. DA tissue levels in sham-operated groups. A, B, day 7, C,D, day 28. A. Striatum.** There was no effect on DA tissue content compared to intact striatum 7 days after sham-operation (sham) ($t=1.90$, $p=0.09$, $df=10$). **B. SN.** There was no effect on DA tissue content compared to intact SN 7 days after sham-operation (sham) ($t=0.86$, $p=0.41$, $df=8$). **C. Striatum.** DA tissue content decreased compared to intact striatum 28 days after sham-operation (sham) ($t=2.78$, $^{*}p=0.03$, $df=7$). **D. SN.** There was no effect on DA tissue content to intact striatum 7 days after sham-operation (sham) ($t=0.67$, $p=0.53$, $df=6$).

**A**

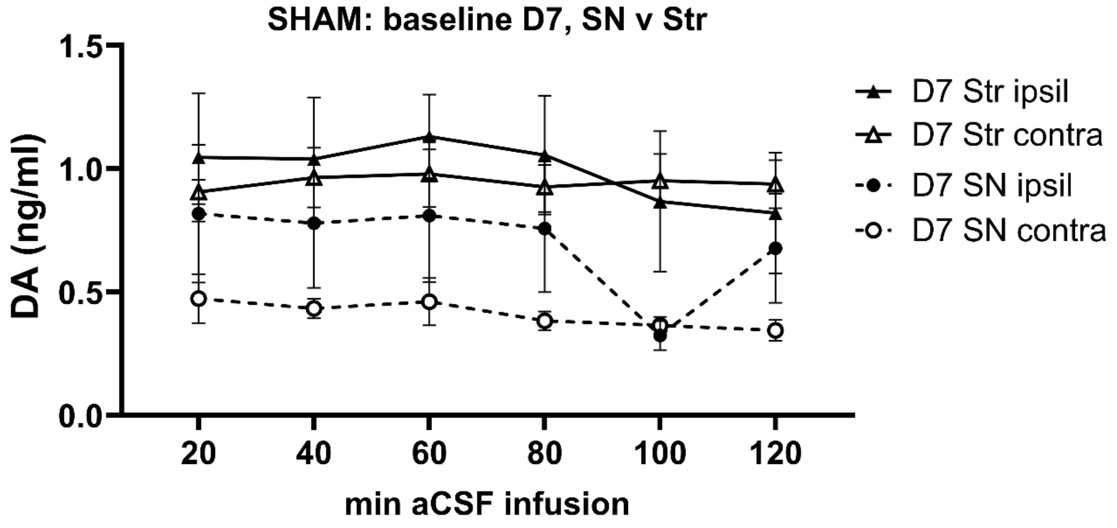


**B.**

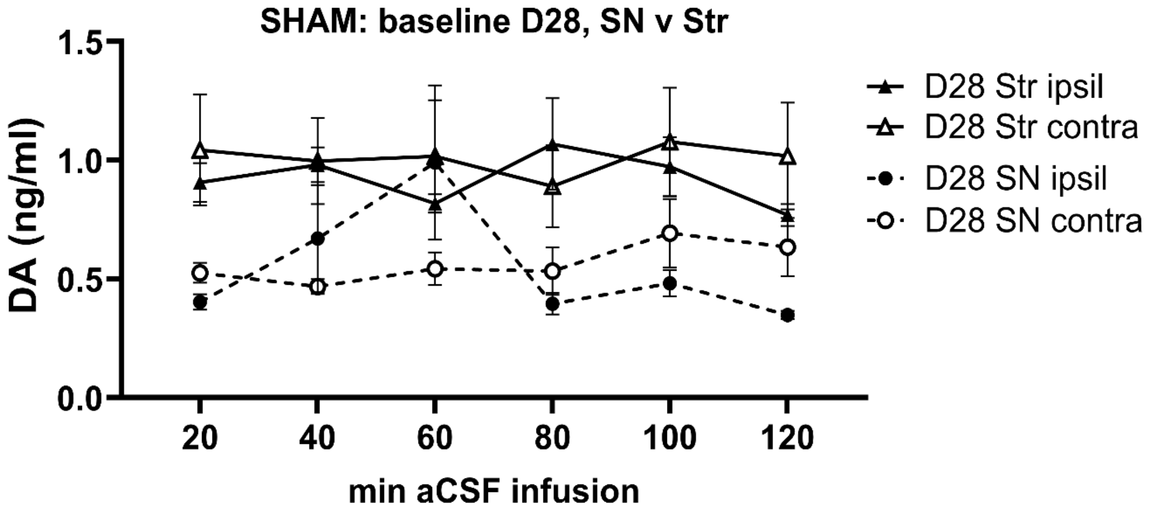


**Suppl Figure 5. Baseline extracellular DA levels sham-operated groups. A. Day 7.** Extracellular DA levels were significantly different between SN and striatum at 7 days after sham-operation, with side of cannulation being no factor. **B. SN.** Extracellular DA levels were significantly different between SN and striatum 28 days after sham-operation, with side of cannulation being no factor.

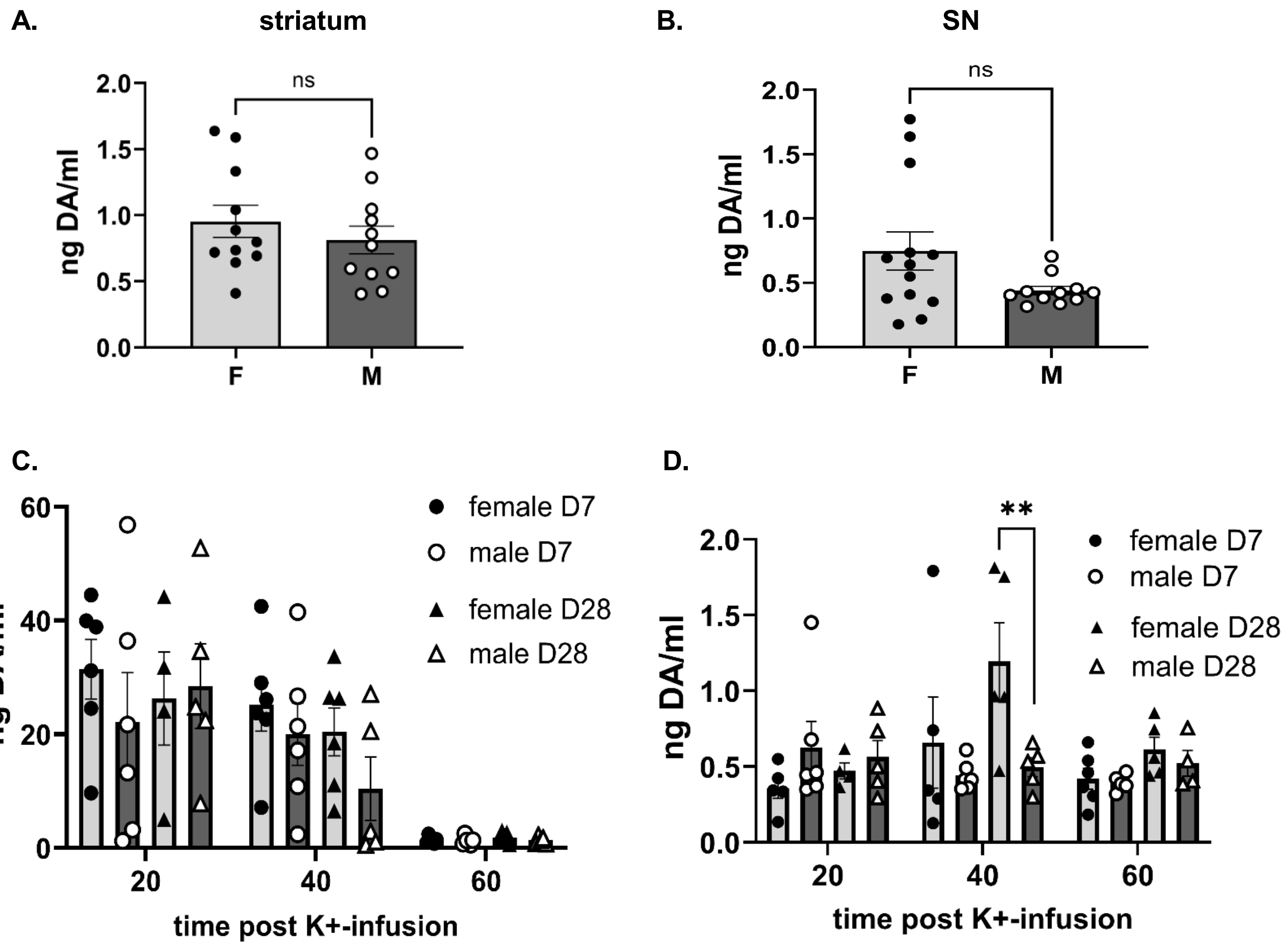


**Suppl Figure 6. Comparison of female v male extracellular DA at baseline (A,B) and K+-stimulated (C,D) conditions in striatum (A,C) and SN (B,D) in sham-operation group. T**ime post-sham operation and time of dialysate collection were not significant variables affecting extracellular baseline DA levels in striatum or SN. Mean extracellular DA levels were compared between (female (F) and male (M) rats in striatum (A) and the SN (B). **A. striatum, baseline** No significant sex-related differences in extracellular DA were observed (t=0.878, *p*=0.39, df=20). **B. SN, baseline.** No significant sex-related differences in extracellular DA were observed in SN (t=1.88, *p*=0.074, df=22), although a trend toward greater levels in females is noted. **C. striatum, K+-infusion.** There were no significant sex-related differences in extracellular DA following K+-infusion in striatum. Although no significant differences in time post sham-operation were observed, results are presented to show results between sexes at day (D) 7 and day 28. **D. SN, striatal K+-infusion.** Although sex differences were not significant across all time points after K+-infusion, there was a significant interaction between time after K+-infusion and sex revealed by 3-way ANOVA and 2-way ANOVA after collapsing days post sham-operation. Collapsing the days together indicated this sex difference was seen 40 min after K+-infusion. Further analysis showed this effect was due to greater extracellular DA in the females only in the D28 cohort (one-way ANOVA, $F(5,22)=4.26$, $p=0.007$), 20 min ($t=0.47$, $p=0.96$, df=22), 40 min ($t=3.75$, $^{**}p=0.003$, df=22), 60 min ($t=0.46$, $p=0.95$ df=22), and not the D7 cohort (one-way ANOVA, $F(5,26)=0.68$, $p=0.65$), 40 min ($t=1.24$, $p=0.67$, df=26).

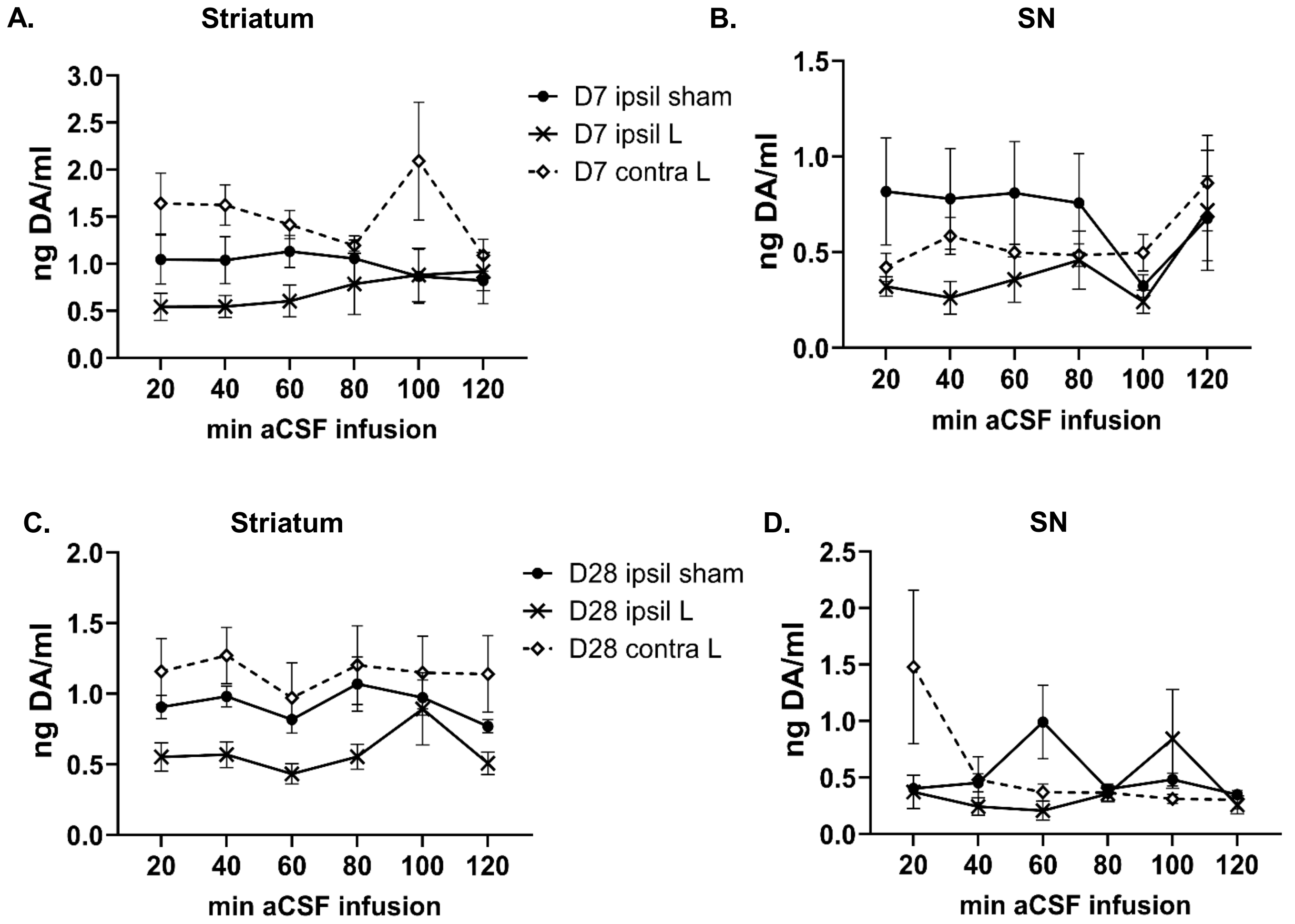


**Suppl Figure 7. Baseline extracellular DA, contralateral to lesioned nigrostriatal pathway.** The following differences were observed in baseline extracellular DA levels with confirmed >70% DA tissue and TH protein loss. **A. Striatum, Day 7.** There was a greater level of extracellular DA contralateral to lesion as compared to striatum ipsilateral to sham (q=4.0, *p*=0.017, df=69) and ipsilateral to lesion (q=6.7, *p*<0.0001, df=69). **B. SN, Day 7.** There was significantly less extracellular DA ipsilateral to lesion as compared to SN ipsilateral to sham (q=3.9, *p*=0.019, df=67) and ipsilateral to lesion (q=6.0, *p*=0.0002, df=73). **C. Striatum, Day 28.** There was a greater level of extracellular DA contralateral to lesion as compared to striatum ipsilateral to sham (q=3.3, *p*=0.057, df=69) and ipsilateral to lesion (q=8.1, *p*<0.0001, df=69). **D. SN, Day 28.** There were no differences in extracellular DA among the 3 groups. Tukey’s multiple comparison test.

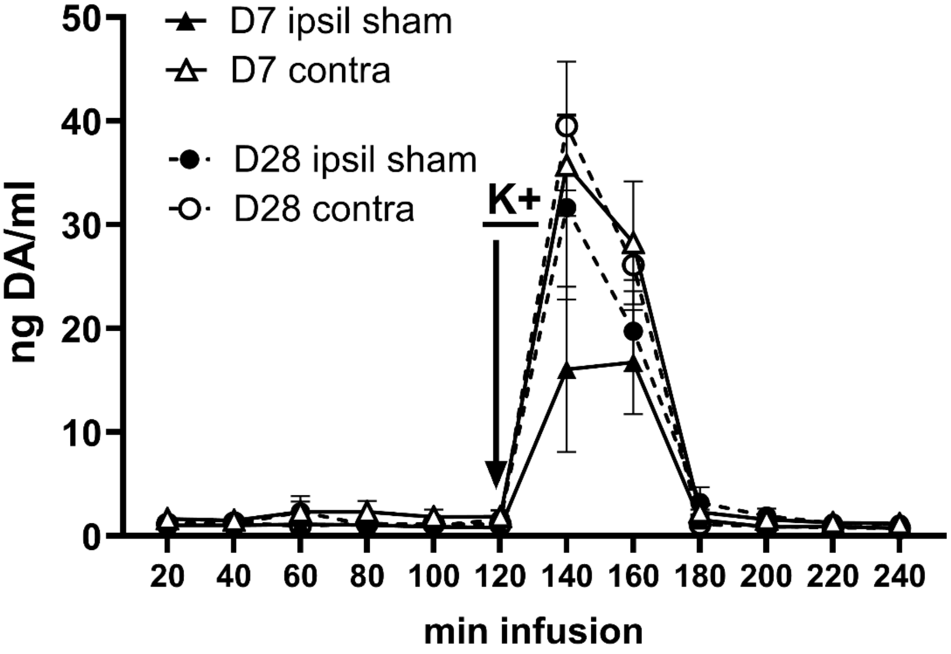


**Suppl Figure 8. K+-evoked extracellular DA, contralateral to lesioned nigrostriatal pathway vs. sham-operation group.** Extracellular DA levels were greater in the striatum contralateral to lesion compared to sham-operated side of the sham-operated groups, particularly at day 7 (D7) ($F(1,16)=4.2$, $p=0.057$).
